# B.1.1.7 and B.1.351 variants are highly virulent in K18-ACE2 transgenic mice and show different pathogenic patterns from early SARS-CoV-2 strains

**DOI:** 10.1101/2021.06.05.447221

**Authors:** Peter Radvak, Hyung Joon Kwon, Martina Kosikova, Uriel Ortega-Rodriguez, Ruoxuan Xiang, Je-Nie Phue, Rong-Fong Shen, James Rozzelle, Neeraj Kapoor, Taylor Rabara, Jeff Fairman, Hang Xie

## Abstract

SARS-CoV-2 continues to circulate globally resulting in emergence of several variants of concern (VOC), including B.1.1.7 and B.1.351 that show increased transmissibility and enhanced resistance to antibody neutralization. In a K18-hACE2 transgenic mouse model, we demonstrate that Both B.1.1.7 and B.1.351 are 100 times more lethal than the original SARS-CoV-2 bearing 614D. Mice infected with B.1.1.7 and B.1.351 exhibited more severe lesions in internal organs than those infected with early SARS-CoV-2 strains bearing 614D or 614G. Infection of B.1.1.7 and B.1.351 also results in distinct tissue-specific cytokine signatures, significant D-dimer depositions in vital organs and less pulmonary hypoxia signaling before death as compared to the mice infected with early SARS-CoV-2 strains. However, K18-hACE2 mice with the pre-existing immunity from prior infection or immunization were resistant to the lethal reinfection of B.1.1.7 or B.1.351, despite having reduced neutralization titers against these VOC. Our study reveals distinguishing pathogenic patterns of B.1.1.7 and B.1.351 variants from those early SARS-CoV-2 strains in K18-hACE2 mice, which will help to inform potential medical interventions for combatting COVID-19.

## Introduction

In December 2019, a highly contagious novel severe acute respiratory syndrome coronavirus 2 (SARS-CoV-2) was transmitted quietly in a local seafood market of Wuhan, Hubei Province, China^1^. From there it has quickly spread around the world through international travelers resulting in the pandemic of coronavirus disease 2019 (COVID-19). Depending on age, sex and comorbidities, humans infected with SARS-CoV-2 experience variable disease severity, ranging from asymptomatic infection to pulmonary dysfunction, multi-organ failure and death^1^. As of June 2, 2021, the COVID-19 pandemic has caused over 170 million confirmed cases and more than 3 million deaths worldwide (https://covid19.who.int/).

SARS-CoV-2 is non-segmented positive-sense, single-stranded RNA virus of bat origin belonging to *Betacoronavirus* family^2^. Its surface glycoprotein – spike (S) contains a receptor binding domain (RBD) that has high affinity to human angiotensin converting enzyme 2 (hACE2) expressed on cell surface and thus allows virus to attach to and fuse with host cell membrane^2–4^. Like other RNA viruses, SARS-CoV-2 undergoes frequent recombination and evolutionary adaption^5^. Soon after the COVID-19 outbreak, a SARS-CoV-2 variant bearing D614G mutation in S outside of the RBD emerged and quickly became the dominant strain in circulation^6, 7^. The D614G mutation enables S protein bind more efficiently to hACE2 and is associated with enhanced virus replication and transmissibility in humans and animal models^6, 7^. As SARS-CoV-2 continues to circulate globally, more SARS-CoV-2 variants have emerged including B.1.1.7 and B.1.351 lineages that were first detected in the UK and South Africa, respectively^8, 9^. The viruses of these two lineages contain multiple mutations in S in addition to widespread D614G^10^. One of the mutations shared by both B.1.1.7 and B.1.351 variants is N501Y in RBD. N501 interacts with several residues of hACE2 including forming a hydrogen bond with Y41 of hACE2 to stabilize the virus-binding hotspot K353 at the RBD-hACE2 interface^3^. The N501Y mutation results in insertion of the aromatic ring of Y501 into a cavity between Y41 and K353 of hACE2, which increases RBD binding affinity to hACE2^11^. This ultimately leads to greater virus transmissibility, as the B.1.1.7 lineage is reportedly 43-90% more transmissible in the UK and 40-50% more transmissible in the US than early SARS-CoV-2 strains^8, 12^. The B.1.351 variant is also estimated to be 50% more transmissible according to the preliminary modeling^9^. E484K is another RBD mutation detected in the B.1.351 lineage and in some of B.1.1.7 strains^10^. E484K confers resistance to several monoclonal antibodies and is able to afford immune escape^13–15^. Pseudoviruses bearing E484K alone or in combination of E484K/N501Y, K417N/E484K or K417N/E484K/N501Y show more resistance to antibody neutralization than the parent pseudoviruses^13–16^. Convalescent or post-vaccination sera are found to less efficiently neutralize B.1.351 variant bearing K417N/E484K/N501Y than the original SARS-CoV-2 strain^15^. In addition to these RBD mutations, both B.1.1.7 and B.1.351 variants also contain several deletions in S protein, including Δ69–70 and Δ144/145 in B.1.1.7 and Δ242–244 in B.1.351^10^. These deletions are found to disrupt an immunodominant epitope in the N-terminal domain of S and alter the antigenicity of S, which not only results in resistance to monoclonal antibodies but also may accelerate SARS-CoV-2 adaptive evolution^17^. Because of increased transmissibility^8, 9, 12^ and enhanced neutralization resistance documented^13–17^, both B.1.1.7 and B.1.351 are classified as variants of concern (VOC) (https://www.cdc.gov/coronavirus/2019-ncov/variants/variant-info.html#Concern). However, the risks of increased disease severity caused by these two variants remain unclear due to many confounding factors such as different geographics, population backgrounds, public health infrastructures (https://www.who.int/publications/m/item/weekly-epidemiological-update-on-covid-19---31-march-2021).

K18-hACE2 transgenic mice was originally developed for the investigations of SARS-CoV-1 pathogenesis^18^. Unlike another hACE2 knock-in mouse strain that does not exhibit mortality following SARS-CoV-2 exposure^6^, this K18-hACE2 mouse strain is susceptible to SARS-CoV-2 and produce various disease manifestations (e.g. lung inflammation and dysfunction, and death) that mimic severe SARS-CoV-2 infection in humans^19^. In this study, we assessed the pathogenicity of B.1.1.7 and B.1.351 variants in K18-hACE2 and compared to that of early SARS-CoV-2 strains containing 614D or 614G. Our results showed that B.1.1.7 and B.1.351 variants not only were more lethal than early SARS-CoV-2 strains but also produced completely different pathogenic profiles in K18-hACE2 mice.

## Results

### CA (B.1.1.7) and SA (B.1.351) are more virulent than early SARS-CoV-2 strains in K18-hACE2 mice

Hemizygous K18-hACE2 mice (confirmed by genotyping) were infected intranasally at 8-10 weeks old for assessing the pathogenicity of SARS-CoV-2 and variants, including (1) USA-WA1/2020 (WA) of lineage A bearing 614D, (2) New York-PV09158/2020 (NY) of lineage B.1.3 bearing 614G, (3) USA/CA_CDC_5574/2020 (CA) of lineage B.1.1.7 and (4) hCoV-19/South Africa/KRISP-EC-K005321/2020 (SA) of lineage B.1.351. Unless otherwise specified, both female and male K18-hACE2 mice were used at approximately 1:1 ratio per infection. Clinical development of diseases was monitored for 14 days post infection (dpi). As early as 5 dpi, K18-hACE2 mice inoculated with 10^2^ 50% tissue culture infectious dose (TCID_50_) per mouse of CA (B.1.1.7) or SA (B.1.351) began to show signs of infection, e.g. becoming motionless, losing body weight (BW) (Fig. 1a), developing hyperthermia (Fig. 1b) and quickly reaching a moribund state (Fig. 1c). By 7 dpi, all K18-hACE2 mice infected with CA (B.1.1.7) died (median survival time: 6 dpi) after losing 15% BW and experiencing severe hyperthermia (Fig. 1a-1c). Mice infected with SA (B.1.351) also lost up to 30% BW (Fig. 1a), exhibited lower body temperature (though statistically insignificant vs naïve mice, Fig. 1b) and all succumbed to death by 10 dpi (median survival time: 7.5 dpi, Fig. 1a and 1c). Intranasal inoculation of 10^2^ TCID_50_/mouse of NY (614G) only resulted in 50% mortality (all in male mice) with no hyperthermia nor overall BW loss (median survival time: 11 dpi, Fig. 1a and 1c). However, K18-hACE2 mice receiving a 10-fold higher inoculation of NY (614G) (10^3^ TCID_50_/mouse) all died within 7 dpi after approximately >15% BW loss (median survival time: 7 dpi, Fig. 1a and 1c). In contrast, intranasal inoculation of WA (614D) at ≥10^3^ TCID_50_/mouse caused no BW drop or hyperthermia and only 1 (male) out of 7 mice died from the infectious dose of 10^4^ TCID_50_/mouse which was 100-fold of CA (1.1.7) or SA (1.1.351) inoculation (Fig. 1a-1c). In subsequent investigations, we thus focused on the pathogenic differences induced by 10^4^ TCID_50_/mouse of WA (614D), 10^3^ TCID_50_/mouse of NY (614G), and 10^2^ TCID_50_/mouse of CA (B.1.1.7) or SA (B.1.351), unless otherwise specified.

**Figure 1.**
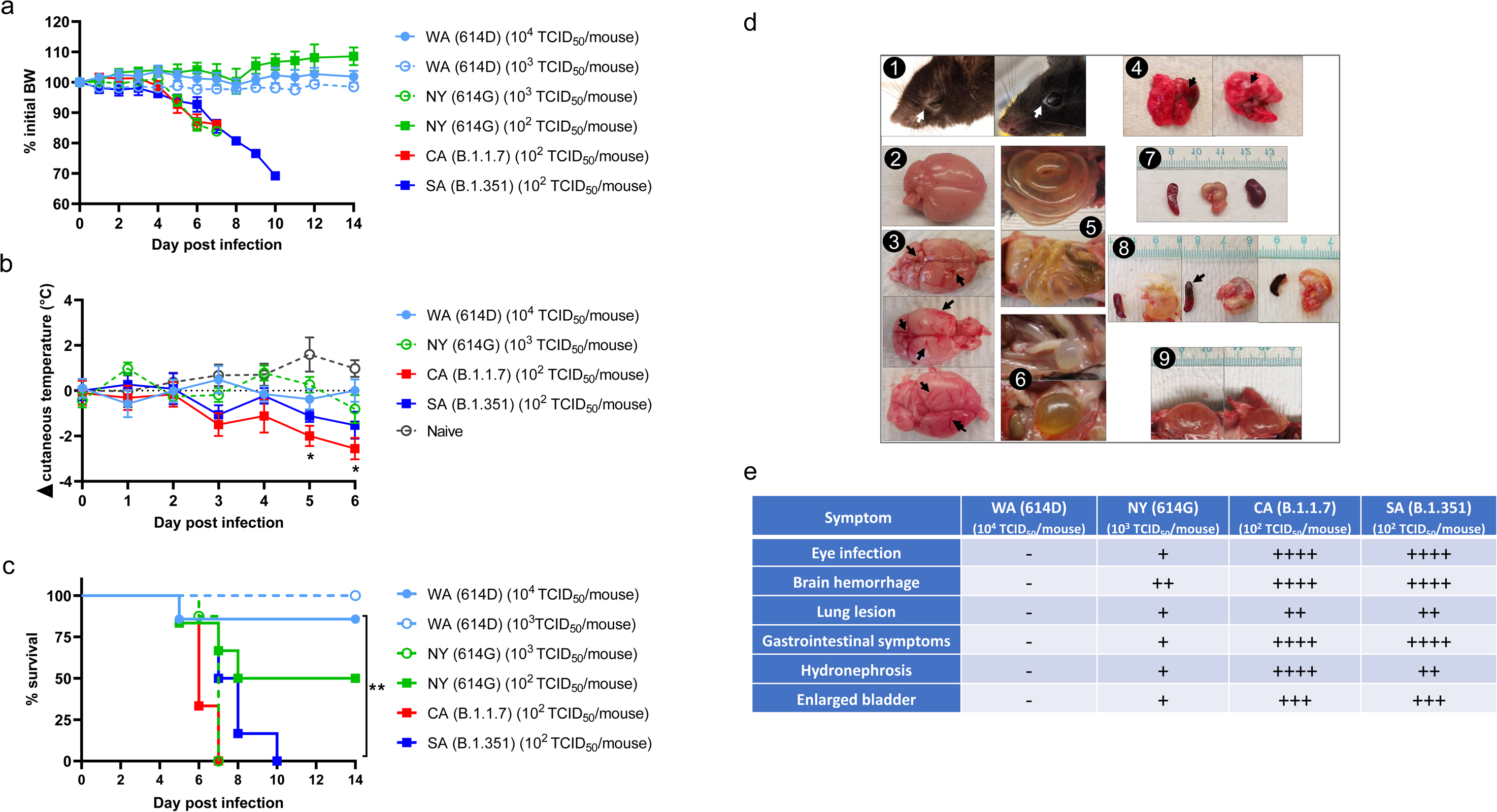
Morbidity, mortality and clinical symptoms of K18-hACE2 mice after infections of SARS-CoV-2 and variants. K18-hACE2 mice of both sexes (approximately 1:1 ratio) were infected intranasally with USA-WA1/2020 (WA) of lineage A bearing 614D, New York-PV09158/2020 (NY) of lineage B.1.3 bearing 614G, USA/CA_CDC_5574/2020 (CA) of lineage B.1.1.7 or hCoV-19/South Africa/KRISP-EC-K005321/2020 (SA) of lineage B.1.351 at indicated doses in an ABSL-3 biocontainment. (a) Body weight (BW); (b) cutaneous temperature change; (c) survival; (d) representative images and (e) frequencies of clinical symptoms after infections. Each group contains 6-8 mice/infection. Data are expressed as mean ± s.e.m. \**p* < 0.05 and \*\**p* < 0.01 by two-way mixed ANOVA (cutaneous temperature change) or by Log-rank (Mantel-Cox) test (survival curves). Images d2 and d7 are from naïve mice and the rest are from SARS-CoV-2 variant infected mice. Arrows indicate the lesions.

In addition to BW loss and hyperthermia (mice do not produce fever), nearly all K18-hACE2 mice inoculated with CA (B.1.1.7) or SA (B.1.351) had eye infection, especially in the corner of eye (Fig. 1d1). Necropsy revealed that intranasal infection of CA (B.1.1.7) or SA (B.1.351) resulted in severe damages in internal organs of K18-hACE2 mice, including hemorrhage in brain (Fig. 1d3 vs naïve in 1d2) and lung (Fig. 1d4), lesions in intestine (Fig. 1d5), bladder (Fig. 1d6), stomach, spleen and kidney (Fig. 1d8 and 1d9 vs naïve in 1d7). These clinical symptoms and lesions were more frequently observed in mice inoculated with CA (B.1.1.7) or SA (B.1.351) than those infected with a 10-fold higher dose of NY (614G) or a 100-fold higher dose of WA (614D) (Fig. 1e). We did not observe sex-specific clinical manifestations in K18-hACE2 mice infected with early SARS-CoV-2 strains. However, female K18-hACE2 mice were found to more likely suffer severe hydronephrosis than male mice following exposure to CA (B.1.1.7) or SA (B.1.351) variant.

We then collected various tissues for viral loads. For WA (614D) or NY (614G) infected mice, tissues were collected after animals were humanely euthanized at 3 dpi and 6 dpi. For CA (B.1.1.7) or SA (B.1.351) infection, tissues were collected at 3 dpi and 5-7 dpi, because the time of infected mice progressed to a moribund state was unpredictable and the quantities of available K18-hACE2 mice were low at the time of this study. After 3 dpi, viral RNA was ready to be detected in brain, lung, liver, spleen, kidney and reproductive organ (ovary in female and seminal vesicles in male) of K18-hACE2 mice (Fig. 2). The expression of hACE2 was confirmed in these organs supporting localized SARS-CoV-2 infection (Supplementary Fig. S1). We monitored viral loads in both testis and seminal vesicles of infected male mice and found that only seminal vesicles consistently had viral RNAs above the detection limit after SARS-CoV-2 infections. Thus, only viral loads in seminal vesicles of infected male mice were reported unless otherwise specified. Higher viral loads were detected in lung and brain homogenates than other organs across all four virus strains (Fig. 2). By 6 dpi, most mice infected with 10^4^ TCID_50_/mouse of WA (614D) had tissue viral RNA levels dropped to the baseline except in lung and brain (Fig. 2). In contrast, mice inoculated with 10-fold less NY (614D) and 100-fold less CA (B.1.1.7) or SA (B.1.351) continued to show elevated viral loads in all organs at 5-7 dpi (Fig. 2), indicating active virus replications in these organs. Interestingly, only a few of those mice infected with CA (B.1.1.7) or SA (B.1.351) had detectable serum IgM ELISA titers specific for viral nucleocapsid (N), spike or RBD at 5-7 dpi (Supplementary Fig. S2). In contrast, nearly all WA (614D) or NY (614G) infected mice had virus-specific IgM ELISA titers ready to be detected in blood as early as 3 dpi (Supplementary Fig. S2).

**Figure 2.**
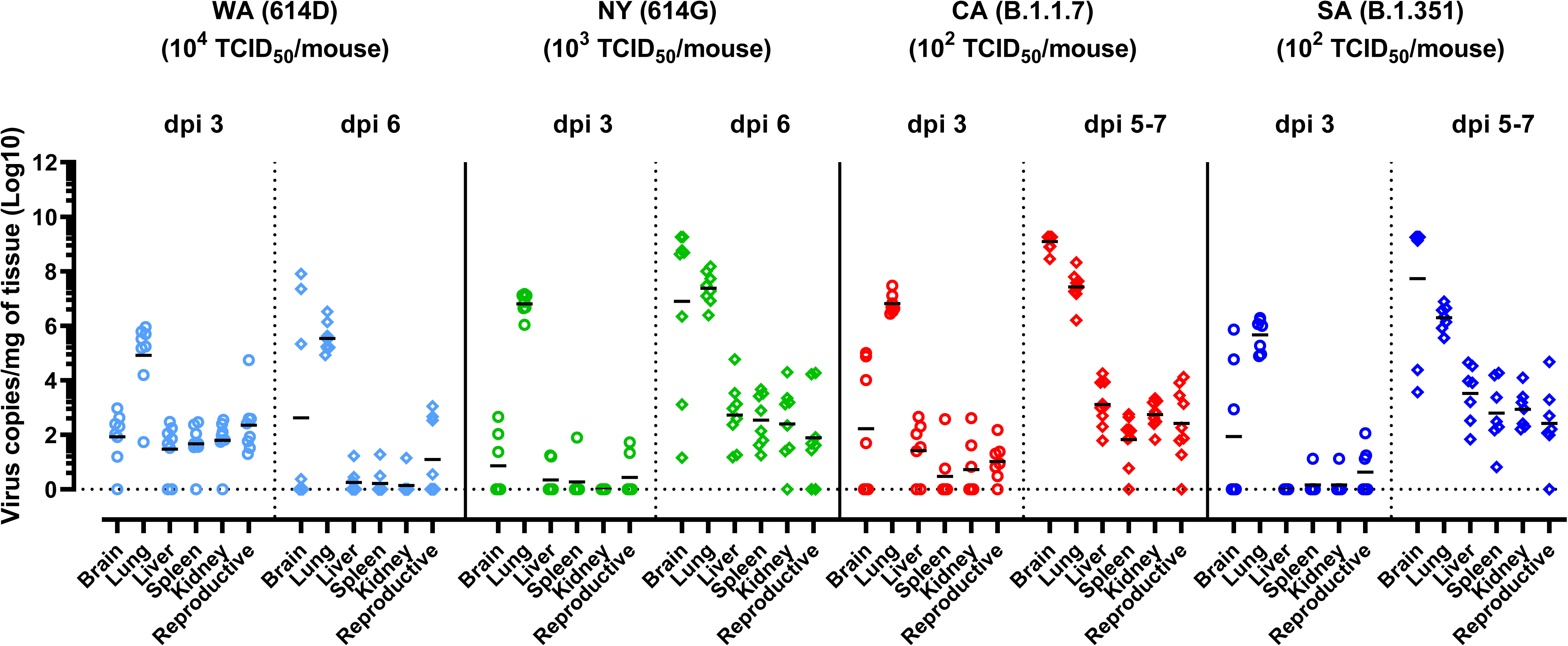
Viral burden in various organs of K18-hACE2 mice after infections of SARS-CoV-2 and variants. K18-hACE2 mice of both sexes (approximately 1:1 ratio) were infected intranasally with USA-WA1/2020 (WA) of lineage A bearing 614D, New York-PV09158/2020 (NY) of lineage B.1.3 bearing 614G, USA/CA_CDC_5574/2020 (CA) of lineage B.1.1.7 or hCoV-19/South Africa/KRISP-EC-K005321/2020 (SA) of lineage B.1.351 at indicated doses in an ABSL-3 biocontainment. Viral RNA copies in brain, lung, liver, spleen, kidney and reproductive organs (ovary or seminal vesicles) at indicated days post infection (dpi) were measured by RT-qPCR (n=7-9 mice/time point/group). Results combining two independent experiments are shown. Short lines represent geometric means and dotted horizontal line indicates the limit of detection.

The results in clinical symptoms, viral burden, morbidity and mortality are consistent, suggesting that both CA (B.1.1.7) and SA (B.1.351) variants are 100-fold more virulent in K18-hACE2 transgenic mice, followed by NY (614G) which is 10-fold more virulent, than WA (614D) that was first detected in the US in the beginning of the COVID-19 pandemic.

### Distinct cytokine profiles are induced in K18-hACE2 mice after infections of SARS-CoV2 and variants

Cytokine storm syndrome has been suggested as a underlying mechanism for respiratory failure in COVID-19 patients^20^. We thus measured proinflammatory cytokines and chemokines in lung and liver homogenates of K18-hACE2 mice at 3 dpi and 5-7 dpi of SARS-CoV-2 and variants (Fig. 3a-3n and Supplementary Fig. S3). Intranasal infections of different SARS-CoV-2 variants resulted in distinct arrays of cytokines secreted by K18-hACE2 mice (Fig. 3a-3n and Supplementary Fig. S3). Mice infected with early SARS-CoV-2 strains tended to produce more chemokines (e.g. IL-13, MCP-1, MIP-1α/β, MIP-2, etc.) in lung and liver than those infected with CA (B.1.1.7) and SA (B.1.351) (Fig. 3a-3n). In contrast, the exposure to CA (B.1.1.7) or SA (B.1.351) variant resulted in higher secretion of type I IFNs, IL-6 and TNF-α in lung and/or liver than those induced by a 10-fold higher infectious dose of NY (614G) or a 100-fold higher infectious dose of WA (614D) (Fig. 3a-3n).

**Figure 3.**
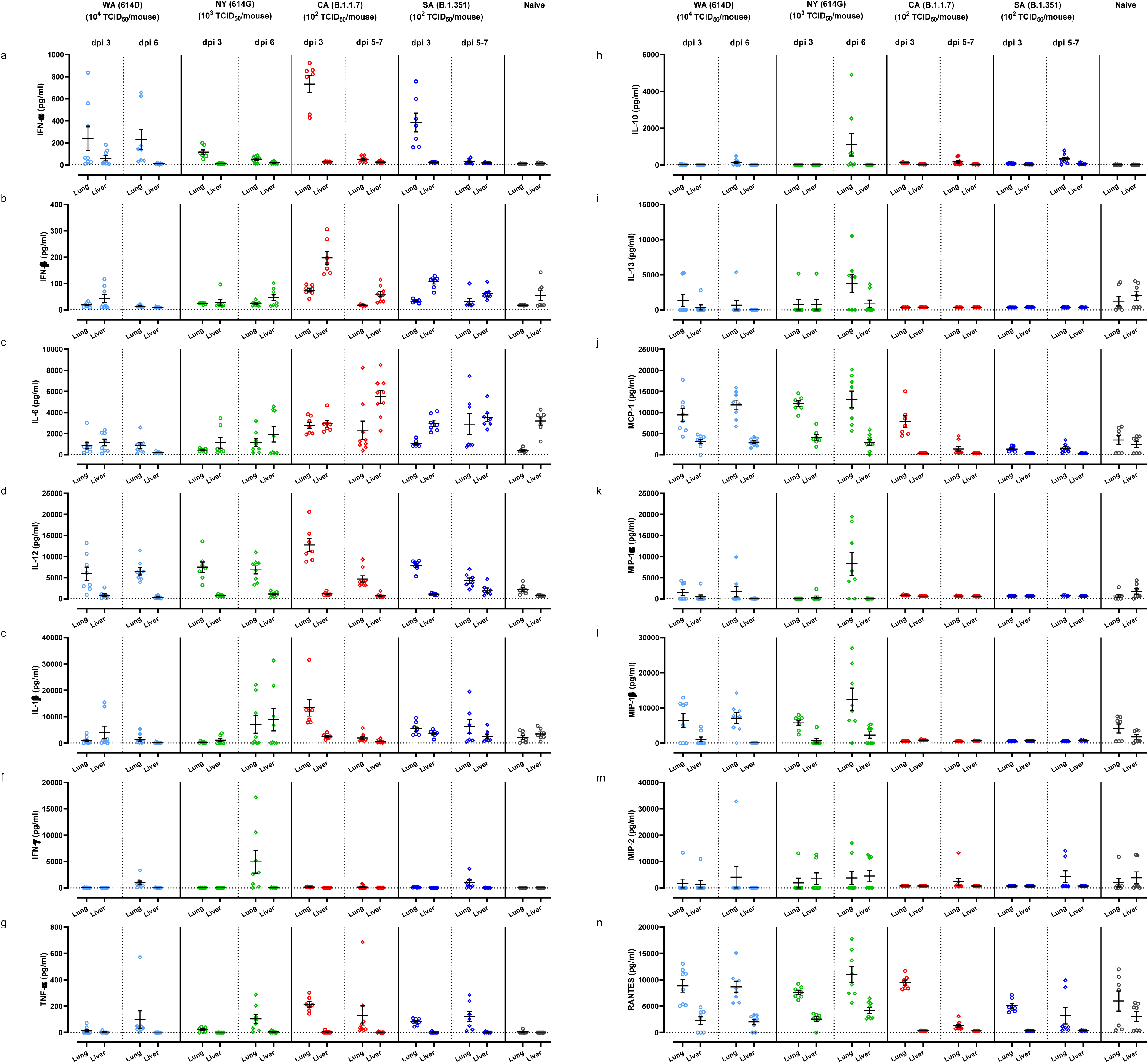
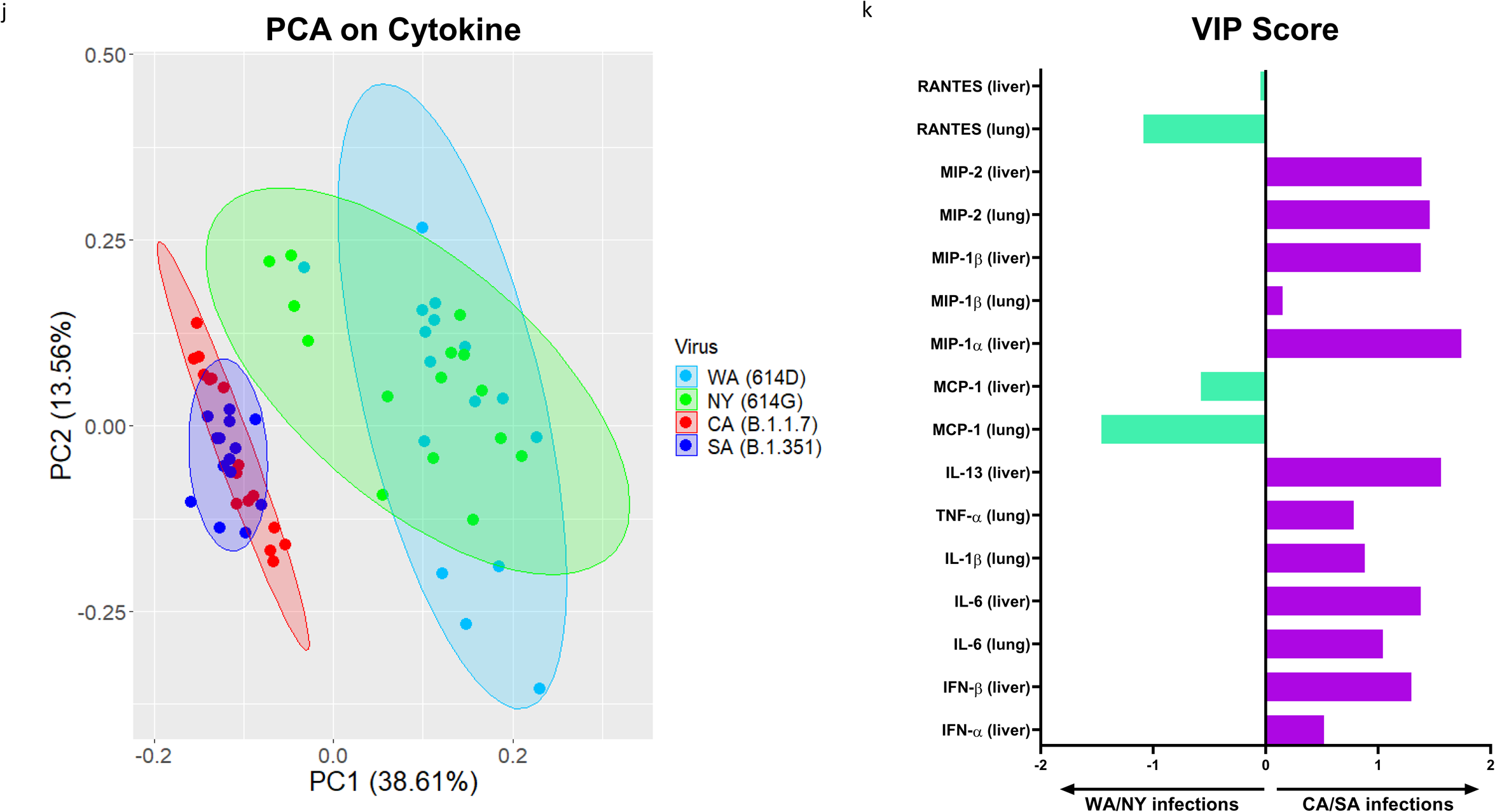
Proinflammatory cytokine profile induced by infections of SARS-CoV-2 or variants. K18-hACE2 mice of both sexes (approximately 1:1 ratio) were infected intranasally with USA-WA1/2020 (WA) of lineage A bearing 614D, New York-PV09158/2020 (NY) of lineage B.1.3 bearing 614G, USA/CA_CDC_5574/2020 (CA) of lineage B.1.1.7 or hCoV-19/South Africa/KRISP-EC-K005321/2020 (SA) of lineage B.1.351 at indicated doses in an ABSL-3 biocontainment. Cytokines in lung and liver homogenates harvested at indicated days post infection (dpi) were measured (n=7-9 mice/time point/group), including (a) IFN-α; (b) IFN-β; (c) IL-6; (d) IL-12; (e) IL-1β; (f) IFN-γ; (g) TNF-α; (h) IL-10; (i) IL-13; (j) MCP-1; (k) MIP-1α; (l) MIP-1β; (m) MIP-2; and (n) RANTES. Results combining two independent experiments are shown. Data are expressed as mean ± s.e.m. Dotted horizontal lines indicate the limit of detection. (j) Principle components analysis (PCA) of all cytokines shown in 3a-3n and Supplementary Figure S3. The most significant two principle components (PC1 and PC2) contain >50% of the variance are shown, which separates WA (614D)/NY (614G) infected mice and CA (B.1.1.7)/SA (B.1.351) infected mice into two groups based on the cytokine profiles; (k) The Variable Importance in Projection (VIP) scores for the cytokine signatures driving the differences between WA/NY and CA/SA infected mice. Only the prime cytokines that were statistically significant are shown. Cytokines pointing towards the left are enriched in WA/NY infected mice, while those pointing right are enriched in CA/SA infected mice. Bar length corresponds to relative importance.

We then performed Principle Component Analysis (PCA) on all cytokines measured in lung and liver homogenates (Fig. 3a-3n and Supplementary Fig. S3) through dimensionality-reduction to extract features that are conserved by individual infections. The most significant two principle components (PC1+PC2 explains more than 50% of the total variation in the cytokine data) grouped WA (614D) and NY (614G) infected mice together while clustered CA (B.1.1.7) and SA (B.1.351) infected mice together (Fig. 3j). This indicated that K18-hACE2 mice infected with CA (B.1.1.7) and SA (B.1.351) variants shared similar cytokine patterns, which were different from the cytokine profiles of mice infected with early WA (614D) and NY (614G) strains. We then conducted Variable Importance in Projection (VIP) to identify the signature cytokine(s) induced by WA/NY or CA/SA infections. The cytokines that were statistically significant between WA/NY and CA/SA groups are shown in Fig. 3k. For infection of CA (B.1.1.7) or SA (B.1.351) variant, multiple cytokines and chemokines in lung and liver were critical to differentiate from those induced by early WA (614D) or NY (614G) strain, including pulmonary MIP-2 (VIP score 1.46), IL-6 (VIP score 1.04), IL-1β (VIP score 0.88), TNF-α (VIP score 0.78) and MIP-1β (VIP score 0.15), and hepatic MIP-1α (1.74), IL-13 (VIP score 1.56), MIP-2 (VIP score 1.39), IL-6 (VIP score 1.38), IFN-β (VIP score 1.30) and IFN-α (VIP score 0.52). However, in mice infected with WA (614D) or NY (614G), only MCP-1 and RANTES secretions in lung (VIP scores: 1.46 for MCP-1 and 1.09 for RANTES) and liver (VIP scores: 0.58 for MCP-1 and 0.046 for RANTES) were important to distinguish from those infected with CA (B.1.1.7) and SA (B.1.351) variants (Fig. 3k).

The above analyses suggest that infection of CA (B.1.1.7) or SA (B.1.351) variant induces similar pulmonary and hepatic cytokine profiles in K18-hACE2 mice, which is distinguishing from those secreted by mice infected with early WA (614D) or NY (614G) strain.

### Pulmonary hypoxia signaling pathway is activated in K18-hACE2 mice infected with NY (614G)

COVID-19 patients often suffer low blood oxygen and “silent hypoxia” has been reported in some severe cases during the early pandemic outbreak^21^. We thus assessed the pulmonary genes involving in hypoxia signaling pathway by qPCR. Substantial upregulations or downregulations of hypoxic genes were found in 3 dpi lung homogenates of K18-hACE2 mice relative to naïve controls (Fig. 4 and Supplementary Fig. S3). The most noticeable changes occurred in the lungs of mice infected with NY (614G) and the expressions of numerous hypoxia-related genes were significantly upregulated (e.g. Anxa2, F3, Hk2, Pim1, Trp53, Txnip, Vdac1 and Vegfa) or downregulated (e.g. Atr, Car9, Expo, Hif3a, Hnf4a, MGDC, etc.) (Fig 4 and Supplementary Fig. S3b). Interestingly, mice infected with CA (B.1.1.7) or SA (B.1.351) variant had considerably less hypoxic genes show altered pulmonary expression at 3 dpi, despite a lethal inoculation administered (Fig 4 and Supplementary Fig. S3c and S3d). Mice administered a high dose of WA (10^4^ TCID_50_/mouse) showed minimum changes in pulmonary expression of hypoxic genes (Fig. 4 and Supplementary Fig. S3a).

**Figure 4.**
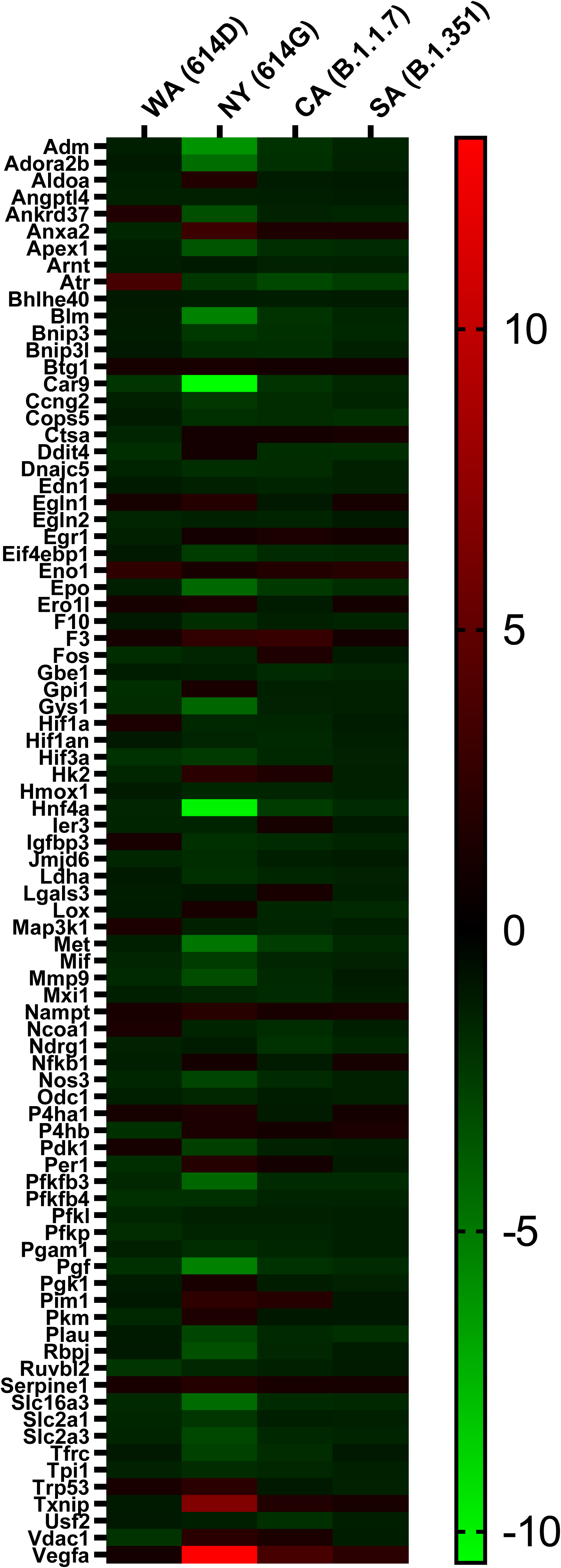
Heatmap of transcriptional genes of the hypoxia pathway following SARS-CoV-2 infections. K18-hACE2 mice of both sexes (approximately 1:1 ratio) were infected intranasally with USA-WA1/2020 (WA) of lineage A bearing 614D (10^4^ TCID_50_/mouse), New York-PV09158/2020 (NY) of lineage B.1.3 bearing 614G (10^3^ TCID_50_/mouse), USA/CA_CDC_5574/2020 (CA) of lineage B.1.1.7 (10^2^ TCID_50_/mouse) or hCoV-19/South Africa/KRISP-EC-K005321/2020 (SA) of lineage B.1.351 (10^2^ TCID_50_/mouse). Hypoxia signaling pathway PCR array was performed using RNA extracted from lung homogenates of naïve or infected mice at day 3 post infection (n= 4 mice/group). Gene regulation as fold changes over naïve mice was used to construct the heatmap.

Nevertheless, only K18-hACE2 mice infected with NY (614G) showed extensive pulmonary hypoxia signaling before reaching to a moribund state; whereas exposure to a lethal dose of CA (B.1.1.7) or SA (B.1.351) or a 10-fold higher dose of WA (614D) resulted in moderate or minimum pulmonary hypoxic pathway activation.

### Infection of CA (B.1.1.7) or SA (B.1.351) variant results in significant elevation of D-dimer in lung, liver, and brain of K18-hACE2 mice

Severely ill COVID-19 patients often develop complications such as pulmonary embolism (PE) and/or deep vein thrombosis (DVT) ^22^. Elevated D-dimer has been associated with increased disease severity and mortality in hospitalized COVID-19 patients^23^. We thus measured tissue-specific D-dimer in lung, liver and brain of K1-hACE2 mice infected with SARS-CoV-2 and variants. Intranasal infection of CA (B.1.1.7) at 10^2^ TCID_50_/mouse resulted in significant elevations of D-dimer in lungs and brains (at a lesser extent) as early as 3 dpi and the levels of D-dimer continued to rise in tissues including livers harvested at later time points (Fig. 5a). This indicated a systemic blood clotting problem developed in K18-hACE2 mice following infection of CA (B.1.1.7). Intranasal infection of SA (B.1.351) also resulted in significant increases of D-dimer in lungs and brains (but not livers) harvested at 3 dpi, which lasted through 5-7 dpi (Fig. 5a). In contrast, K1-hACE2 mice infected with early WA (614D) or NY (614G) strain showed no significant elevation of D-dimer in tissues harvested, except slight rises of hepatic D-dimer level as compared to naïve mice (Fig. 5a). Interestingly, brain D-dimer levels showed no correlation with viral loads detected in brains (Pearson correlation coefficient r=0.022); whereas pulmonary and hepatic D-dimer levels correlated well with viral loads in individual tissues – r=0.344 for lung (*p*= 0.0041) and r= 0.341 for liver (*p*= 0.0044), respectively (Fig. 5b). However, brain D-dimer levels strongly correlated with D-dimer levels detected in lung (r= 0.929, *p*< 0.00001) and liver (r= 0.417, *p*< 0.001) as well (Fig. 5b). The PCA on tissue-specific D-dimer responses yielded two distinct groups (PC1+PC2>98%), which mice infected with early WA (614D) or NY (614G) were clustered together; whereas mice infected with CA (B.1.1.7) or SA (B.1.351) variant were together in a separate group (Fig. 5c). Moreover, VIP scores for D-dimer in lung, liver and brain were 1.17, 0.61 and 1.12, respectively, all pointing to the cluster of CA/SA infections (Fig. 5d).

**Figure 5.**
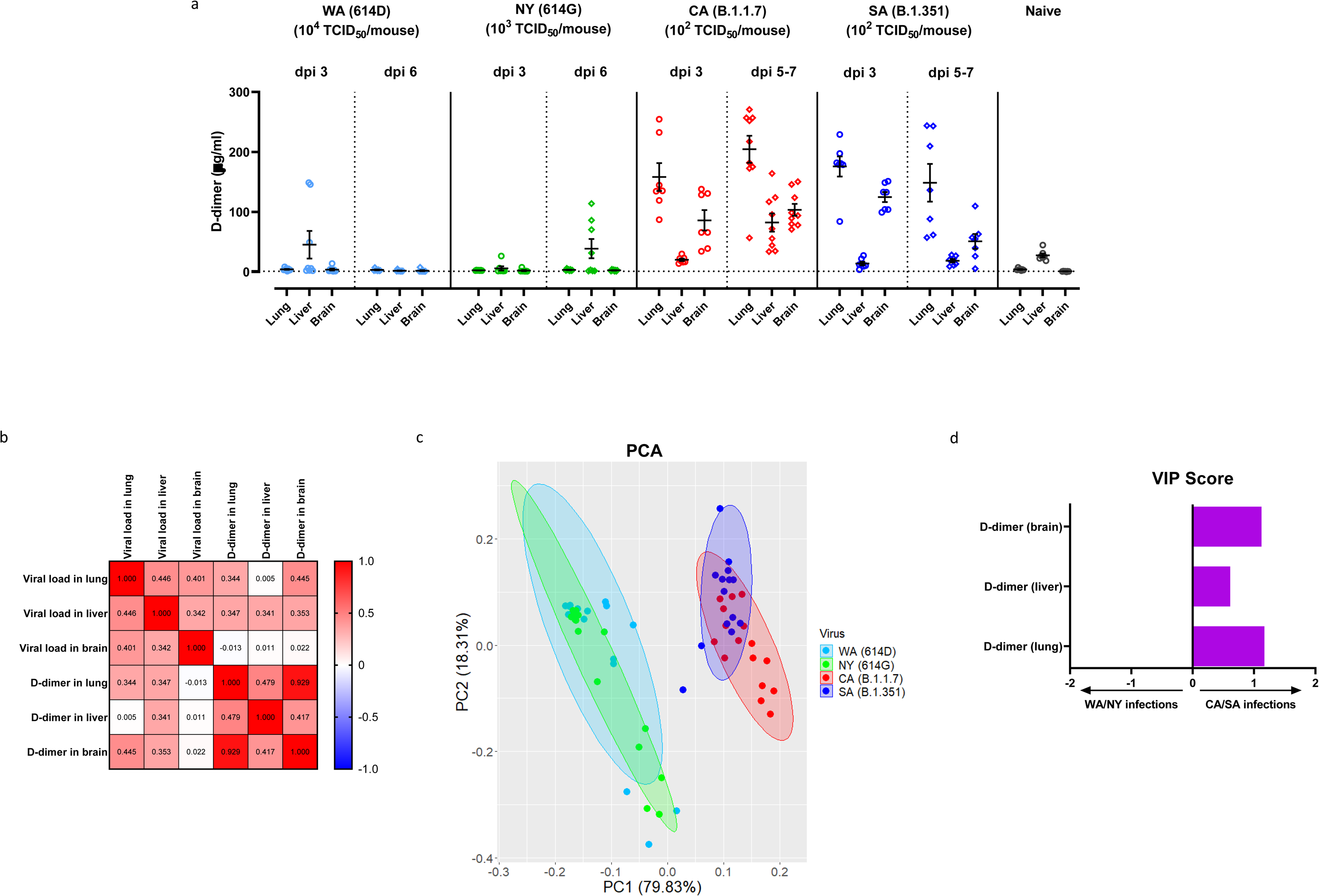
D-dimer levels in various tissues following infections of SARS-CoV-2 and variants. K18-hACE2 mice of both sexes (approximately 1:1 ratio) were infected intranasally with USA-WA1/2020 (WA) of lineage A bearing 614D, New York-PV09158/2020 (NY) of lineage B.1.3 bearing 614G, USA/CA_CDC_5574/2020 (CA) of lineage B.1.1.7 or hCoV-19/South Africa/KRISP-EC-K005321/2020 (SA) of lineage B.1.351 at indicated doses in an ABSL-3 biocontainment. (a) D-dimer levels in lung, liver and brain homogenates at indicated days post infection (dpi). Results combining two independent experiments are shown. Data are expressed as mean ± s.e.m (n=7-9 mice/time point/tissue). Dotted horizontal lines indicate the limit of detection. (b) Correlation matrix of tissue-specific D-dimer levels with viral loads by Pearson correlation analysis after the log transformation of the data. Individual Pearson correlation coefficient r values are shown; (c) Principle components analysis (PCA) of D-dimer levels shown in Figure 5a. The two most significant principle components (PC1+ PC2> 98%) separate WA (614D)/NY (614G) infected mice and CA (B.1.1.7)/SA (B.1.351) infected mice into two groups based on D-dimer levels; (d) The Variable Importance in Projection (VIP) scores for the D-dimer levels in lung, liver and brain that drive the differences between WA/NY and CA/SA infected mice. Bar length corresponds to relative importance.

These results suggest tissue-specific D-dimer levels in lung, brain and liver (at a lesser extent) were critical to distinguish infection of CA (B.1.1.7) or SA (B.1.351) variant from those infected with early WA (614D) or NY (614G) strain in K18-hACE2 mice. That D-dimer levels in brain, lung and liver strongly correlated, suggest the blood clotting problems observed in these organs were related to the same systemic cause.

### K18-hACE2 mice with prior SARS-CoV-2 infection or immunization are protected from a lethal infection of CA (B.1.1.7) or SA (B.1.351) variant

A critical question remaining to be answered is whether COVID-19 patients who have recovered from early infections develop effective immunity to protect them from later exposure to SARS-CoV-2 variants such as CA (B.1.1.7) or SA (B.1.351). To address this, we reinfected K18-hACE2 that had survived from prior infections of WA (614D) or NY (614G) in Fig. 1a-1c with 10^2^ TCID_50_/mouse of CA (B.1.1.7) or SA (B.1.351) variant. These mice had developed >3 log10 IgG ELISA titers specific for RBD or S from prior infections (Fig. 6a). The sera collected right before reinfection also had >3 log10 microneutralization (MN) titers against live WA (614D) and NY (614G) viruses, but exhibited 38-46% reduction in MN titers against live CA (B.1.1.7) (*p*< 0.0001 vs WA or *p*< 0.001 vs NY, Fig. 6b) or 32-40% reduced MN titers against live SA (B.1.351) (*p*< 0.0001 vs WA or *p*< 0.01 vs NY, Fig. 6b). Despite having significantly reduced MN antibody titers against new variants, these mice all survived a lethal reinfection of CA (B.1.1.7) or SA (B.1.351) with no obvious BW drops (Fig. 6c and 6d). K18-hACE2 mice immunized with RBD or S emulsified with AddaS03™ (an AS03-like research grade adjuvant) also developed good antigen-specific IgG ELISA titers (Fig. 6e). Immunized mice also had markedly reduced MN titers against the variants especially SA (B.1.351) as compared to the titers toward WA (614D) (Fig. 6f). Nevertheless, all mice immunized with adjuvanted S were protected from lethal challenges of NY (614G) or SA (B.1.351) without showing substantial morbidity (Fig. 6g and 6h). Immunization of adjuvanted RBD also resulted in 75% survival (3 out of 4 mice/group/virus) from the lethal NY (614G) or SA (B.1.351) challenge (Fig. 6g and 6h).

**Figure 6.**
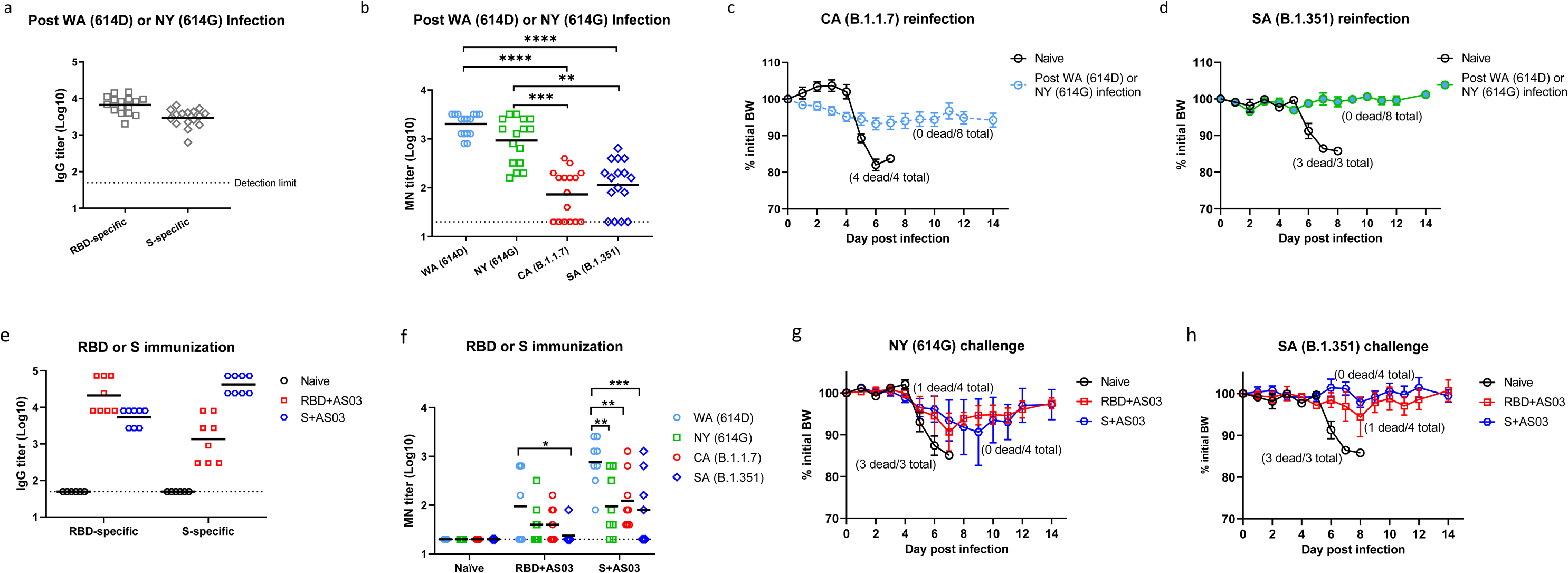
Antibody responses, morbidity and mortality following reinfection or challenge of SARS-CoV-2 SA variant. K18-hACE2 mice that survived from prior infection of USA-WA1/2020 (WA) of lineage A bearing 614D or New York-PV09158/2020 (NY) of lineage B.1.3 bearing 614G were reinfected at 6-8 weeks later with USA/CA_CDC_5574/2020 (CA) of lineage B.1.1.7 or hCoV-19/South Africa/KRISP-EC-K005321/2020 (SA) of lineage B.1.351 at the dose of 10^2^ TCID_50_/mouse in an ABSL-3 biocontainment. Right before reinfection, mouse sera were analyzed for (a) IgG titers specific for spike (S) or receptor binding domain (RBD) by ELISA or (b) microneutralization (MN) titers for SARS-CoV-2 and variants (n= 16 mice/group). Body weight (BW) and survival of mice from prior infection (n= 8 mice/group) and naïve K18-hACE2 mice (n= 3 mice/group) were monitored for two weeks after reinfection of CA (c) or SA (d) variant. Separate sets of K18-hACE2 mice were immunized with RBD or S emulsified with AS03-like adjuvant and were challenged with 10^3^ TCID_50_/mouse of NY (614G) or 10^2^ TCID_50_/mouse of SA (B.1.351). (e) Serum IgG ELISA titers specific for RBD or S or (f) MN titers for SARS-CoV-2 and variants before challenge (n= 8 mice/group). BW and survival of immunized mice (n= 4 mice/group) and naïve K18-hACE2 mice (n= 3 mice/group) were monitored for two weeks after challenge of NY (g) or SA (h) variant. Short lines represent geometric means and dotted horizontal lines indicate the limit of detection. BW are expressed as mean ± s.e.m. \**p* < 0.05, \*\**p* < 0.01, \*\*\**p* < 0.001 and **** *p* < 0.0001 by one-way nonparametric ANOVA with Dunn’s multiple comparisons test after log-transformation of MN titers. (d) and (h) were conducted at the same time with the same naïve mice as the control.

These results together suggest the SARS-CoV-2-specific immunity developed from prior infection or immunization is effective in protecting K18-hACE2 mice from later exposure to CA (B.1.1.7) or SA (B.1.351) variant, despite showing up to 46% reduction in antibody neutralizing capacities toward these variants.

## Discussion

B.1.1.7 and B.1.351 variants contain N501Y in RBD in addition to widespread D614G in S protein^10^. Both N501Y and D614G have been shown to increase RBD binding to hACE2 and promote virus entry and replication in humans and animal models^6, 7^. The N501Y alone or in combination with K417N, E484K or K417N/E484K of S mutations identified in B.1.351 variant also greatly enhances viral infectivity in HEK293T with hACE2 overexpression^16^. Additionally, the deletion of H69/V70 in S1 subunit is reported to increase the infectivity of B.1.1.7 variant and benefit more cleaved S incorporated into virions^24^. In this study, K18-hACE2 mice infected with CA (B.1.1.7) or SA (B.1.351) variant had comparable or higher viral loads in various organs than those inoculated with a 10-fold higher dose of NY (614G) or a 100-fold higher dose of WA (614D). The systemic dissemination of virus in K18-hACE2 transgenic mice was in consistence with tissue tropism of hACE2 expression. Some infected mouse organs had hACE2 expression lower than naïve controls, which may be due to virus-induced hACE2 shedding/downregulation or cell death in these individual tissues^19^. Nevertheless, the overall expression of hACE2 in most tissues was enhanced after SARS-CoV-2 infection.

We observed that male K18-hACE2 mice were more susceptible than female mice to early SARS-CoV-2 strains. This is consistent with the meta-analysis of global cases that shows male sex is a risk factor for COVID-19 and male patients have higher odds for intensive care admission or death during the early pandemic^25^. Sex-specific differences in hormones, immunity and social behaviors may account for the relative resistance of females to COVID-19^26–28^. Additionally, human reproductive organs have high levels of ACE2 expression and SARS-CoV-2 RNA has been detected in testis and semen from male patients, and placenta and vaginal mucous membrane from pregnant women, respectively^29–31^. However, the long-term effects of COVID-19 on both male and female fertility remain unknown. In K18-hACE2 mice, viral RNA was also detected in ovary of female mice or seminal vesicles/testis of male mice following infection of SARS-CoV-2 or variants. Thus, K18-hACE2 mice would be an excellent model to assess the impact of SARS-CoV-2 infection on the fertility of both sexes.

Unlike those early COVID-19 cases showing sex differences, infections with B.1.1.7 or B.1.351 variant were found to have men and women almost in the equal ratio^32^. Human infections of B.1.1.7 or B.1.351 variants also had significantly higher adjusted odds ratios for hospitalization and intensive care admission than non-VOC cases based on the data collected from seven EU/EEA countries^32^. These facts indicate that the infection patterns caused by B.1.1.7 or B.1.351 variant are different from those early SARS-CoV-2 strains. We also observed that B.1.1.7 and B.1.351 variants exhibit at least 100-fold higher lethality than the early WA (614D) strain in K18-hACE2 mice. Compared to WA (614D) infected mice, infection of B.1.1.7 or B.1.351 variants resulted in more severe damages to internal organs, including brain hemorrhage, eye infection, gastrointestinal lesions and other clinical relevant manifestations that mirror the severity of COVID-19 symptoms in humans^29, 33^. Both male and female K18-hACE2 appeared equally susceptible to B.1.1.7 or B.1.351 variant, except that hydronephrosis was more frequently spotted in female mice after exposures.

The analyses of cytokines secreted in the lungs and livers of K18-hACE2 mice revealed that infection of B.1.1.7 and B.1.351 variants resulted in a completely different inflammatory profile from those infected with early SARS-CoV-2 strains bearing 614D or 614G. Both MCP-1 (CCL2) and RANTES (CCL5) secreted in lung and liver rose as the tissues-specific cytokine signatures for mice infected with WA (614D) and NY (614G). MCP-1 and RANTES belong to the C-C chemokine family and are potent chemotactic factors that regulate migration and infiltration of monocytes/macrophages or T cells, respectively. An early study in K18-hACE2 mice has reported that lung infiltration of myeloid cells and activated CD8+ T cells was associated with enhanced pulmonary expression of MCP-1/CCL2 and RANTES/CCL5 induced by WA (614D) infection^19^. In this study, K18-hACE2 infected with B.1.1.7 and B.1.351 variants exhibited more complicated tissue-specific cytokine signatures than those induced by early SARS-CoV-2 strains. Several myeloid cell chemoattractants showed enhanced pulmonary and hepatic secretions in K18-hACE2 mice following exposures to B.1.1.7 or B.1.351 variant, including MIP-1α/CCL3, MIP-1β/CCL4 and MIP-2/CXCL2. Moreover, infection of B.1.1.7 and B.1.351 resulted in significant elevation of several key inflammatory cytokines in K18-hACE2 mice, including TNF-α (lung), IL-1β (lung), IL-6 (lung and liver), IL-13 (liver), IFN-α (liver) and IFN-β (liver). These inflammatory cytokines are also reportedly elevated in critically ill COVID-19 patients^1, 34–40^. Elevated IL-6 in particular is associated with SARS-CoV-2 viral loads in serum and is strongly correlated with the severity of COVID-19 in humans^1, 35, 36, 40, 41^. Patients with severe or critical COVID-19 are reported to have diminished systemic Type I IFN responses but high pulmonary expression of type I and III IFNs^40, 42^. Excessive IFN signaling not only impairs lung repair by inducing protein p53 but also increases host susceptibility to bacterial superinfections in a mouse model of influenza infection^43^. Dysregulated Type I IFN responses have also been linked to SARS-CoV-1 induced lethality in both humans and mice^44, 45^. Thus, the tissue-specific cytokine signatures identified in B.1.1.7 and B.1.351 infected K18-hACE2 mice not only indicate the severity of infection but also provide insights into potential immunomodulatory interventions for combatting COVID-19.

Low blood oxygen is one of the early clinical signs present in COVID-19 patients. Some patients may experience dangerously low oxygen levels without developing dyspnea or showing other signs of respiratory distress– the so-called “silent hypoxia”^46^. Hypoxia has been reported to be independently associated with mortality in hospitalized COVID-19 patients and has been suggested as an indicator for COVID-19-related intensive care admission^47^. In this study, K18-hACE2 mice exhibited extensive hypoxia signaling in lungs at 3 dpi of NY (614G), but had minimum hypoxic genes upregulated or downregulated after WA (614D) infection. The latter may be because a sublethal dose of WA (614D) was used to inoculate K18-hACE2 mice. However, K18-hACE2 mice infected with CA (B.1.1.7) or SA (B.1.351) variant had fewer hypoxic genes altered than those infected with NY (614G), despite they all were administered a lethal infectious dose. Furthermore, infections of CA (B.1.1.7) and SA (B.1.351) variants resulted in significant D-dimer depositions in lungs and brains (livers at less extent) of K18-hACE2 mice as compared to those exposed to early WA (614D) or NY (614G) strain. Pulmonary embolism and venous/arterial thromboembolism are frequently diagnosed in critically ill COVID-19 patients and are significantly associated with disease severity (intensive care admission or death)^22, 48, 49^. Elevated D-dimer indicating blood clot formation has been used as a diagnostic marker for pulmonary embolism in COVID-19 patients, and hospitalized patients with superhigh D-dimers often have poor prognosis and low chance of survival^1, 22, 23, 41, 50^. The deposition of D-dimers in the vital organs along with a lack of extensive pulmonary hypoxia signaling before death indicates that the pathogenic patterns of B.1.1.7 and B.1.351 variants are different from those of early SARS-CoV-2 strains in K18-hACE2 mice. Both pulmonary and hepatic D-dimer levels were also found to correspond with local viral loads, indicating virus replication induced tissues damages in these mice. Interestingly, brain D-dimer levels of infected mice showed no correlation with local viral loads but were strongly associated with pulmonary D-dimer levels. These are likely due to SARS-CoV-2 caused systemic endothelial injury and microthrombosis^49, 51^. The postmortem histopathological examination reveals that many COVID-19 patients suffered multifocal microvascular injury in brains, e.g. congested blood vessels, fibrinogen leakage, and microhemorrhages in addition to widespread thrombosis in pulmonary vessels ^49, 51^. K18-hACE2 mice infected with CA (B.1.1.7) or SA (B.1.351) variant also had severe hemorrhages in brain before they succumbed to sudden death. These differences in pulmonary hypoxia signaling, tissue-specific D-dimer deposition and cytokine file confirm that the B.1.1.7 and B.1.351 variants have distinct pathogenic characteristics from early SARS-CoV-2 strains.

Sera collected from K18-hACE2 with prior exposure to early SARS-CoV-2 strains or immunized with adjuvanted S or RBD had significantly reduced neutralization titers against CA (B.1.1.7) and SA (B.1.351). This is consistent with the reports that viruses bearing escape mutations such as Δ69-70 and Δ144/145 in B.1.1.7 or K417N/E484K/N501Y in B.1.351 are able to avoid recognition by monoclonal antibodies or show resistance to neuralization by convalescent/post-vaccination sera^13–17^. Despite having reduced neutralizing antibodies, K18-hACE2 mice with prior SARS-CoV-2 exposure or immunized with AS03-like adjuvanted S all survived the lethal challenge of CA (B.1.1.7) or SA (B.1.351) variant with minimum BW loss. Immunization of AS03-like adjuvanted RBD also protected the majority of K18-hACE2 mice from the lethality of SA (B.1.351) or NY (614G). Infection or immunization induced cellular immunity may contribute to the protection observed in these K18-hACE2 mice with low neutralization titers against variants. It has been reported that SARS-CoV-2 infection induces long-lasting memory B cells and T cells in humans^52, 53^. Individuals with prior infection are able to develop robust T cell and memory B cell responses against B.1.1.7 and B.1.351 variants after receiving first mRNA vaccine dose^54^. Investigations on characterizing T cell and B cell responses in K18-hACE2 mice following exposure to different SARS-CoV-2 variants or immunization are currently underway.

In summary, the current study in K18-hACE2 mice reveals that B.1.1.7 and B.1.351 variants of concern are at least 100 times more lethal than the original SARS-CoV-2 WA (614D) strain detected in the beginning of the pandemic. The in vivo lethality of B.1.1.7 and B.1.351 variants is companied by BW loss, hyperthermia and severe internal organ damages in infected K18-hACE2 mice. Infection of B.1.1.7 and B.1.351 variants also produces a completely different pathogenic profile from that of early SARS-CoV2 strains (614D or 614G), which is characterized as distinct tissue-specific proinflammatory cytokine signatures, significant D-dimer depositions in vital organs and a lack of extensive pulmonary hypoxia signaling before death. Despite having reduced serum neutralization titers, K18-hACE2 mice with the pre-existing immunity from prior infection or immunization are protected from reinfection of B.1.1.7 or B.1.351 variant. The current study not only identifies the in vivo pathogenic characteristics of B.1.1.7 and B.1.351 variants but also provide insights into potential medical interventions to control inflammation, mitigate tissue damages and improve survival outcome of COVID-19 patients.

## Methods

### Cells and viruses

Vero E6 (CRL-1586; American Type Culture Collection or ATCC) was maintained in high-glucose Dulbecco’s modified Eagle’s medium (DMEM) supplemented with 10% fetal bovine serum (FBS), 1% penicillin/streptomycin and 10 mM HEPES (pH 7.3) at 37 °C, 5% CO2. The SARS-CoV-2 clinical isolates were obtained through BEI Resources (see Acknowledgements for details), including (1) USA-WA1/2020 (WA) of lineage A bearing 614D (ATCC# NR-52281), (2) New York-PV09158/2020 (NY) of lineage B.1.3 bearing 614G (ATCC# NR-53516), (3) USA/CA_CDC_5574/2020 (CA) of lineage B.1.1.7 (ATCC# NR-54011) and (4) hCoV-19/South Africa/KRISP-EC-K005321/2020 (SA) of lineage B.1.351 (ATCC# NR-54008). Infectious stocks were prepared by inoculating 90% confluent Vero E6 cells in DMEM supplemented with 3% FBS at 37 °C, 5% CO2 until observation of cytopathic effect. Supernatants were centrifuged and passaged through a 0.45-μm filter. Aliquots were stored in a secured −80 °C freezer until use. All work with live SARS-CoV-2 isolates was performed in an animal biosafety level (ABSL) 3 laboratory equipped with advanced access control devices and by personnel equipped with powered air-purifying respirators.

### Determination of 50% tissue culture infectious dose (TCID50)

Virus infectivity was determined using an ELISA-based TCID_50_ method adopted from a protocol used for titrating influenza virus^55^. Briefly, virus was diluted in ½ log10 in minimum essential medium (MEM) supplemented with 2% aseptically filtered bovine serum album (BSA) solution (Millipore-Sigma). Serially diluted virus was added to Vero E6 cells (2X10^4^ cells/well in MEM with 2% BSA) pre-seeded in 96-well tissue culture plates. Each virus dilution was tested in octuplets. After incubation at 37 °C, 5% CO2 for two days, virus-infected cells were detected using an in-house rabbit polyclonal antibody specific for SARS-CoV-2 N/M/E followed by horseradish peroxidase-conjugated goat anti-mouse IgG (Invitrogen). TCID_50_ was calculated using the Reed-Muench method.

### Recombinant proteins

The DNA sequences encoding the full-length S (residues 1-1213) and RBD (residues 319-541) of WA (GenBank: MN985325.1) with a T4 trimerization domain and a 6xHis tag at the C-terminus were codon-optimized and were subcloned into pcDNA5/FRT mammalian expression vector (ThermoFisher). Expi293 Expression system (ThermoFisher) was used to produce recombinant SARS-CoV-2 S and RBD proteins according to the manufacturer’s protocol. The cultured media harvested on day 6 post-transfection were centrifuged at 27,000 × g, 4 °C for 60 min. The supernatants were concentrated and diafiltrated through Amicon Ultra-15 10K (EMD Millipore) in the equilibration buffer (50 mM Tris-HCl, 150 mM NaCl and EDTA-free protease inhibitor cocktail, pH 8.0). The diafiltrated supernatants were then loaded to equilibrated Ni affinity columns (Cytiva, Marlborough, MA) and bound proteins were eluted in the presence of 500 mM imidazole. Expressed S and RBD proteins were purified using an AKTA Pure system (Cytiva) with >90% purity as confirmed by SDS-PAGE. Expression of the full-length E, M and N proteins^56^ was carried out in propriety Xpress CF™ system with reactions in thin-film Petri dish incubated at 25°C overnight without shaking. For recombinant E and M, 0.1% Brij35 was added to the cellfree reactions and purification buffers to aid solubility and slow aggregation. Expressed E, M and N were purified by Ni-NTA affinity chromatography with >85% purity by SDS-PAGE.

### Mice

Hemizygous B6.Cg-Tg(K18-ACE2)2Prlmn/J (K18-hACE) female mice (JAX Stock No. 034860) and noncarrier c57BL/6J male mice (JAX Stock No. 000664) were mated to maintain live colonies at the ABSL2 facility of FDA White Oak Vivarium. Only hACE2 (+) offspring as confirmed by genotyping (Transnetyx) were retained and were used for experiments. At the age of 8-10 weeks, K18-hACE mice of both sexes were randomly assigned to experimental groups at approximately 1:1 ratio and were transferred to the ABSL3 facility for infection. All K18-hACE mice were ear tagged and were housed at two per microcage in the ABSL3 facility. Under isoflurane anesthesia, mice were inoculated with SARS-CoV-2 or variants via intranasal route. Body weight (BW) and mortality were monitored daily for up to 14 days post infection (dpi). Cutaneous temperature was also measured on mouse tail surface using a rodent infrared thermometer (Braintree Scientific) following infection. Mice becoming moribund or reaching humane endpoints (e.g. 30% BW loss) were immediately euthanized. Mice survived from prior infection of WA (614D) or NY (614G) were reinfected at 6-8 weeks later with 10^2^ TCID_50_/mouse of CA (B.1.1.7) or SA (B.1.351). In separated experiments, K18-hACE2 mice of both sexes were primed and boosted at 3 weeks intervals with S (10 µg/dose) or RBD (20 µg/dose) emulsified with AddaS03™ (InvivoGen) via intramuscular route. Sera were collected right before the challenges for antibody determination. All procedures were performed according to the animal study protocols approved by the FDA White Oak Animal Program Animal Care and Use Committee.

### Viral burden and hACE2 expression

Tissues were weighed and homogenized in PBS (pH7.2) (10%, v/w) with ceraminc beads in a Fisherbrand™ Bead Mill 24 Homogenizer (FisherScientific). Total RNA was extracted from tissue homogenates using the RNeasy Plus Mini Kit (Qiagen) and was immediately reverse transcribed and amplified using the High-Capacity cDNA Reverse Transcription Kit (Thermo Fisher Scientific) following the manufacturer’s protocol. Copies of SARS-CoV-2 nucleocapsid (N) gene in homogenized tissues were determined using QuantiNova SYBR Green PCR kit (Qiagen) along with 2019-nCoV RUO Kit (Integrated DNA Technologies) as gene specific primers at the final concentration of 500 nM. Quantitative amplification in Stratagene MX3000p qPCR system (Agilent) was conducted according to the following program: 95 °C for 120 s, 95 °C for 5 s (50 cycles) and 60 °C for 18 s. Threshold cycle (Ct) values were calculated using MxPro qPCR software (Agilent) and the N gene copies in individual tissues were interpolated from a standard curve constructed by serial dilutions of a pCC1-CoV2-F7 plasmid expressing SARS-CoV-2 N^57^. hACE2 expression in different mouse organs was determined similarly using hACE2 gene specific primer set (Integrated DNA Technologies, Assay ID: Hs.PT.58.27645939) at the final concentration of 500 nM. The copies of expressed hACE2 were interpolated from a standard curve created by serial dilutions of a pSBbi-bla ACE2 plasmid expressing hACE2 (kindly provided by J Yewdell, NIAID/NIH). A value of 1 was assigned if gene copies were below the detection limits.

### Hypoxia signaling pathway PCR array

Total RNA extracted from lungs on dpi 3 was reverse transcribed as described above and was mixed with RT^2^ SYBR Green ROX qPCR Mastermix (Qiagen) to perform RT² Profiler™ PCR Array Mouse Hypoxia Signaling Pathway (Qiagen) real-time PCR in Stratagene MX3000p qPCR system under these cycling conditions: hold for 10 min at 95°C, followed by 40 cycles of 15 s at 95°C and 60 s at 60°C. Individual gene expressions based on Ct values were normalized using Gusb as the internal housekeeping gene and were calculated for fold changes using Qiagen web-based GeneGlobe data analysis tool. Heatmap was generated using Prism 8.4.3 (GraphPad, San Diego, CA).

### Cytokine and chemokine detection

Proinflammatory cytokines and chemokines in mouse lung and liver homogenates were determined in a MESO QuickPlex SQ 120 imager using MSD multiplex kits (Meso Scale Diagnostic, Rockville, MD) according to the manufacturer’s protocols. Type I interferons (IFNs) in mouse lung and liver homogenates were quantitated using VeriKine mouse IFN-α or Verikine High Sensitivity mouse IFN-β ELISA kits (PBL Assay Science, Piscataway, NJ).

### D-dimer measurement

Detection of D-dimer in tissue homogenates was carried out using mouse D-Dimer ELISA Kit (MyBioSource) according to the manufacturer’s instructions.

### Antibody ELISA

All mouse sera were heat-inactivated before antibody assessment. ELISA was performed in 96-well microtiter plates pre-coated with 1 µg/ml of recombinant S, RBD or N protein. Bound antibodies were detected using peroxidase-conjugated goat anti-mouse IgM heavy chain (Southern Biotech) or IgG (H+L) antibodies (Invitrogen) followed by One-step TMB substrate (ThermoFisher). Optical density (OD) at 450 nm was measured using a Victor V multilabel reader (PerkinElmer, Waltham, MA). ELISA titers were interpolated based on a standard curve constructed using in-house developed rabbit polyclonal antibodies specific for S, RBD or N, respectively.

### Microneutralization (MN) assay

Neutralizing antibody titers against live SARS-CoV-2 and variants were determined using a cell-based MN assay. Briefly, heat-inactivated mouse sera were serially diluted and were incubated with 100 TCID_50_/well of live virus at room temperature for 1 hour. The virus-serum mixtures were then added to Vero E6 cells (2X10^4^ cells/well) pre-seeded in 96-well tissue culture plates and were incubated at 37°C, 5% CO2 for 2 days. Cells were then fixed with 4% paraformaldehyde for 30 min followed by permeabilization in 0.1% NP-40 detergent for 15 min. Virus-infected cells were detected using in-house raised rabbit anti-S polyclonal antibody along with peroxidase-conjugated goat anti-rabbit IgG (H+L) secondary antibody (Invitrogen). MN titers represented the reciprocal of the highest serum dilution that yielded >50% reduction in OD values as compared to wells containing virus only. A MN titer of 20 was assigned if no neutralization was observed at the initial serum dilution of 1:40.

### Histology

The whole brains and lungs harvested from infected mice were fixed in 10% neutral buffered formaldehyde for at least two weeks before being embedded in paraffin for histology (Histoserv). Tissues embedded in paraffin were sectioned and stained with hematoxylin and eosin by the standard protocols (Histoserv). Uninfected mouse brains and lungs were used as negative controls and stained in parallel. Tissue sections were visualized using a Leica Aperio AT2 slide scanner with Aperio ImageScope DX clinical viewing software (Histoserv).

### Statistical analysis

Multivariate Principal Components Analysis (PCA) and Partial Least Squares Discriminant Analysis (PLSDA) were used to analyze cytokine profile and D-dimer among different infection groups. Principal Component Analysis (PCA) was used to visualize if samples from different virus exposures were separated into discrete groups within the two-dimensional principal components (PC) space. The variable importance in projection (VIP) scores, which measures the variable’s importance, were reported from the PLSDA model. Linear models were also performed to associate tissue-specific D-dimer with viral loads in individual organs. R version 4.0.3 (https://www.r-project.org/) was used to perform all the above analyses. Prism 8.4.3 (GraphPad, San Diego, CA) was used to conduct one-way nonparametric ANOVA or two-way mixed ANOVA after the log transformation of antibody titers, or Log-rank (Mantel-Cox) survival test. A *p* value of <0.05 was considered statistically significant.

### Data availability

Relevant data and reagents are available from the corresponding author upon reasonable request and or by Material Transfer Agreement.

## Acknowledgments

This work was supported by FDA/CBER intramural SARS-CoV-2 pandemic fund. The following reagents were obtained through BEI Resources, NIAID, NIH: SARS-Related Coronavirus 2: (1) Isolate USA-WA1/2020, NR-52281 (deposited by the Centers for Disease Control and Prevention); (2) Isolate New York-PV09158/2020, NR-53516; (3) Isolate USA/CA_CDC_5574/2020, NR-54011 (deposited by the Centers for Disease Control and Prevention); (4) Isolate hCoV-19/South Africa/KRISP-EC-K005321/2020, NR-54008 (contributed by Alex Sigal and Tulio de Oliveira). The authors sincerely appreciate the support of FDA/CBER Biosafety team and White Oak Vivarium staff in this study. The authors thank Drs. Tony Wang and Zhiping Ye (Division of Viral Products, FDA/CBER) for technical and administrative assistances at the beginning of setting up SARS-CoV-2 BSL3. The pSBbi-bla ACE2 plasmid was kindly provided by Dr. Jonathan W. Yewdell (NIAID/NIH).

## AUTHOR CONTRIBUTIONS

H. X. conceived and designed the study. P.R., H.J.K., M.K., U.O-R., and H.X. conducted the experiments, performed biological assays, and analyzed the data. R.X. and H.X. performed the statistical analyses. P.R., J.P., R.S., J.R., N.K., T.R., and J.F. expressed recombinant proteins. H.X., P.R., H.J.K., M.K., U.O-R., J.P., and J.R. composed the manuscript. H.X. modified and revised the final version.

---

**Supplementary Figure S1.**
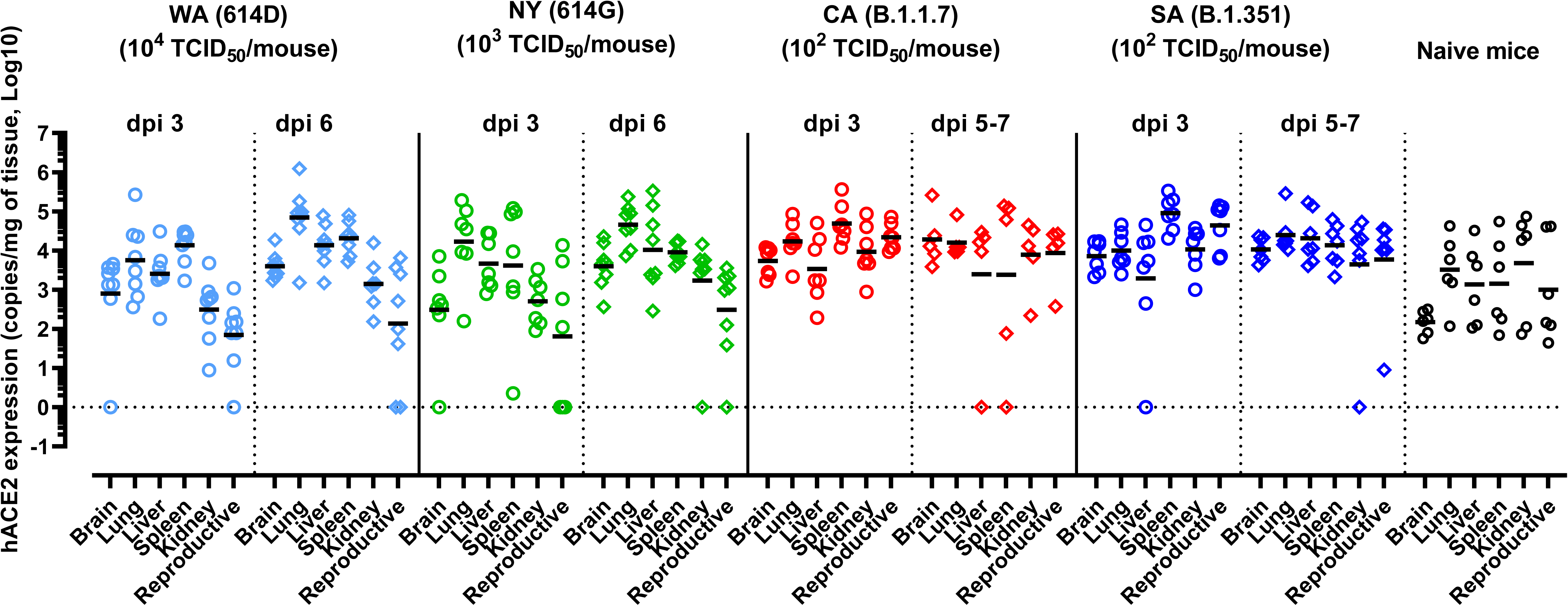
hACE2 expression in various organs of K18-hACE2 mice after infections of SARS-CoV-2 and variants. K18-hACE2 mice of both sexes (approximately 1:1 ratio) were infected intranasally with USA-WA1/2020 (WA) of lineage A bearing 614D (10^4^ TCID_50_/mouse), New York-PV09158/2020 (NY) of lineage B.1.3 bearing 614G (10^3^ TCID_50_/mouse), USA/CA_CDC_5574/2020 (CA) of lineage B.1.1.7 (10^2^ TCID_50_/mouse) or hCoV-19/South Africa/KRISP-EC-K005321/2020 (SA) of lineage B.1.351 (10^2^ TCID_50_/mouse). hACE2 expression in brain, lung, liver, spleen, kidney and reproductive organs (ovary or seminal vesicles) at indicated days post infection (dpi) were measured by RT-qPCR (n=7-9 mice/time point/group). Data are expressed as mean ± s.e.m. Short lines represent geometric means and dotted horizontal line indicates the limit of detection.

**Supplementary Figure S2.**
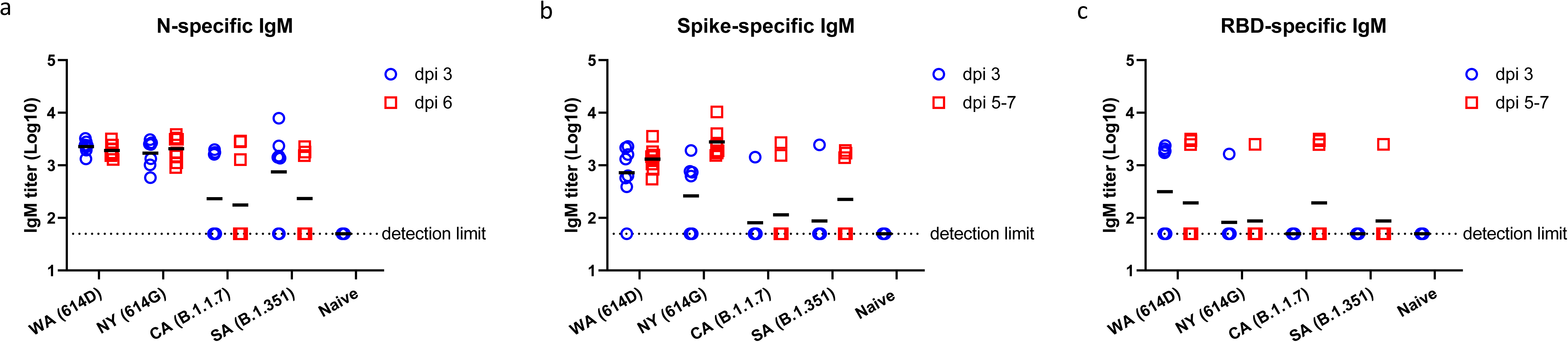
Detection of virus-specific IgM in K18-hACE2 after infections of SARS-CoV-2 and variants. K18-hACE2 mice of both sexes (approximately 1:1 ratio) were infected intranasally with USA-WA1/2020 (WA) of lineage A bearing 614D (10^4^ TCID_50_/mouse), New York-PV09158/2020 (NY) of lineage B.1.3 bearing 614G (10^3^ TCID_50_/mouse), USA/CA_CDC_5574/2020 (CA) of lineage B.1.1.7 (10^2^ TCID_50_/mouse) or hCoV-19/South Africa/KRISP-EC-K005321/2020 (SA) of lineage B.1.351 (10^2^ TCID_50_/mouse). Sera were collected at indicated days post infection (dpi) for IgM ELISA titers specific for SARS-CoV-2 spike, receptor binding domain (RBD) or nucleocapsid (N). Short lines represent geometric means and dotted horizontal lines indicate the limit of detection.

**Supplementary Figure S3.**
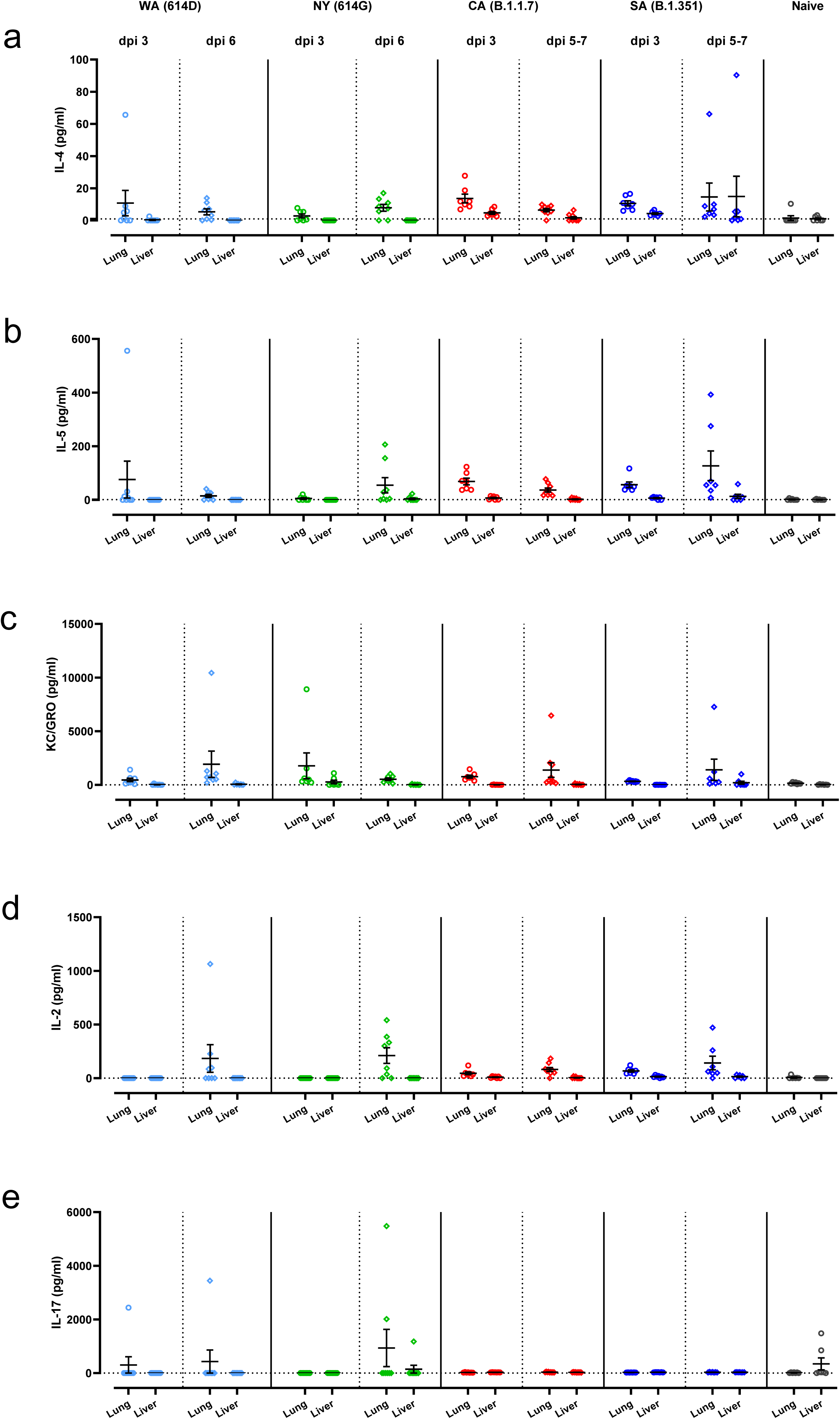
Additional proinflammatory cytokines induced by infections of SARS-CoV-2 or variants. K18-hACE2 mice of both sexes (approximately 1:1 ratio) were infected intranasally with USA-WA1/2020 (WA) of lineage A bearing 614D, New York-PV09158/2020 (NY) of lineage B.1.3 bearing 614G, USA/CA_CDC_5574/2020 (CA) of lineage B.1.1.7 or hCoV-19/South Africa/KRISP-EC-K005321/2020 (SA) of lineage B.1.351 at indicated doses in an ABSL-3 biocontainment. Additional mouse cytokines in lung and liver homogenates harvested at indicated days post infection (dpi) were measured (n=7-9 mice/time point/group), including (a) IL-4; (b) IL-5; (c) KC/GRO; (d) IL-2; and (e) IL-17. Mean ± s.e.m are shown and dotted horizontal lines indicate the limit of detection.

**Supplementary Figure S4.**
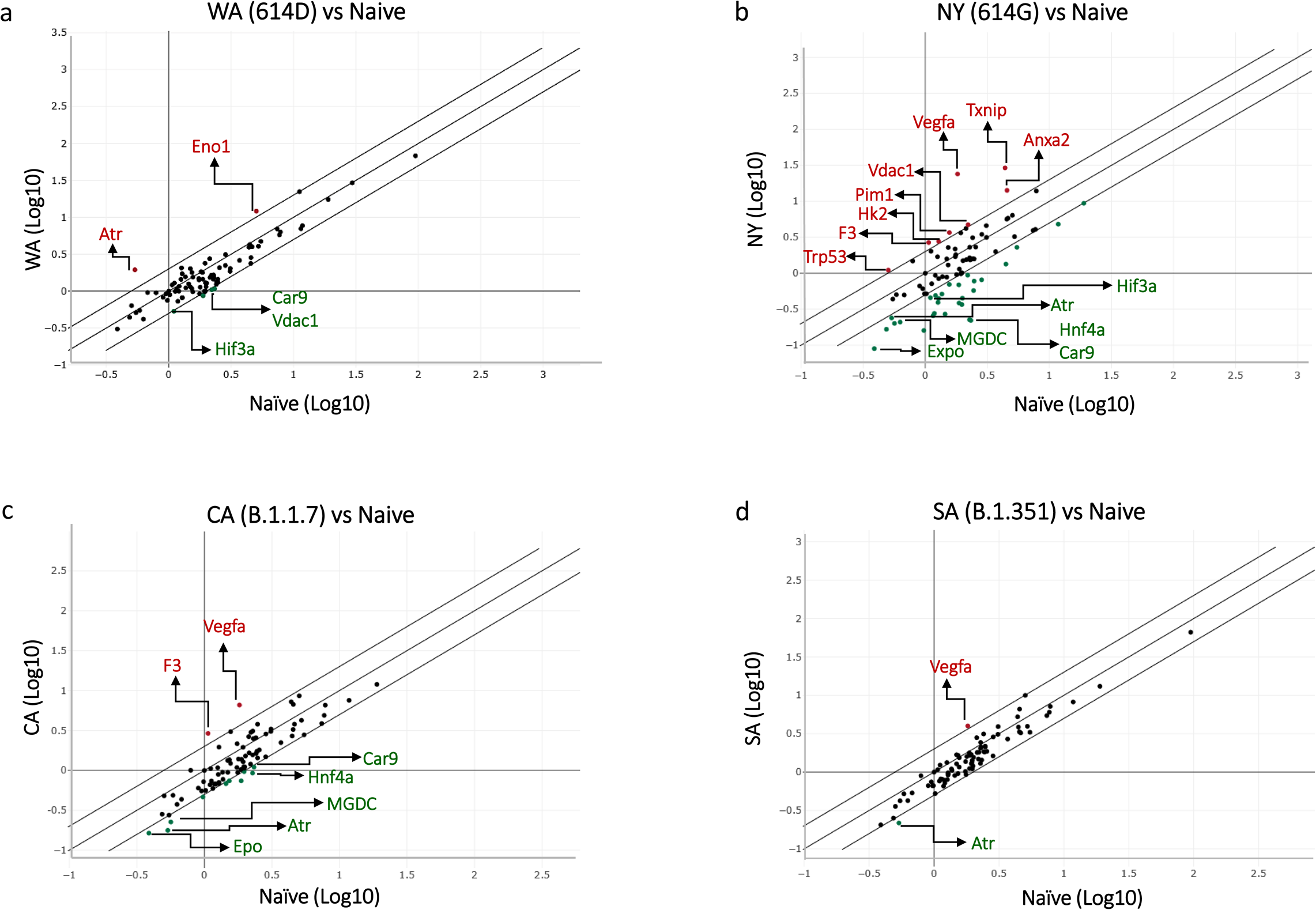
The correlation scatter plot of pulmonary gene expression of hypoxia signaling pathway after SARS-CoV-2 infections. K18-hACE2 mice of both sexes (approximately 1:1 ratio) were infected intranasally with USA-WA1/2020 (WA) of lineage A bearing 614D (10^4^ TCID_50_/mouse), New York-PV09158/2020 (NY) of lineage B.1.3 bearing 614G (10^3^ TCID_50_/mouse), USA/CA_CDC_5574/2020 (CA) of lineage B.1.1.7 (10^2^TCID_50_/mouse) or hCoV-19/South Africa/KRISP-EC-K005321/2020 (SA) of lineage B.1.351 (10^2^ TCID_50_/mouse). Hypoxia signaling pathway PCR array was performed using RNA extracted from lung homogenates of naïve or infected mice at day 3 post infection (n= 4 mice/group). The average fold changes of individual genes vs naïve mice are presented in the correlation scatter plots. Genes beyond the 2-fold change lines are marked by red (upregulation) or green (downregulation) dots. Typical genes that involve in the hypoxia signaling pathway are labeled.

## References

1 Huang, C. et al. Clinical features of patients infected with 2019 novel coronavirus in Wuhan, China. Lancet 395, 497–506, doi:10.1016/S0140-6736(20)30183-5 (2020).

2 Zhou, P. et al. A pneumonia outbreak associated with a new coronavirus of probable bat origin. Nature 579, 270–273, doi:10.1038/s41586-020-2012-7 (2020).

3 Shang, J. et al. Structural basis of receptor recognition by SARS-CoV-2. Nature 581, 221–224, doi:10.1038/s41586-020-2179-y (2020).

4 Lan, J. et al. Structure of the SARS-CoV-2 spike receptor-binding domain bound to the ACE2 receptor. Nature 581, 215–220, doi:10.1038/s41586-020-2180-5 (2020).

5 Gupta, R. et al. SARS-CoV-2 (COVID-19) structural and evolutionary dynamicome: Insights into functional evolution and human genomics. J Biol Chem 295, 11742–11753, doi:10.1074/jbc.RA120.014873 (2020).

6 Zhou, B. et al. SARS-CoV-2 spike D614G change enhances replication and transmission. Nature 592, 122–127, doi:10.1038/s41586-021-03361-1 (2021).

7 Volz, E. et al. Evaluating the Effects of SARS-CoV-2 Spike Mutation D614G on Transmissibility and Pathogenicity. Cell 184, 64–75 e11, doi:10.1016/j.cell.2020.11.020 (2021).

8 Davies, N. G. et al. Estimated transmissibility and impact of SARS-CoV-2 lineage B.1.1.7 in England. Science 372, doi:10.1126/science.abg3055 (2021).

9 Tegally, H. et al. Detection of a SARS-CoV-2 variant of concern in South Africa. Nature 592, 438–443, doi:10.1038/s41586-021-03402-9 (2021).

10 Boehm, E. et al. Novel SARS-CoV-2 variants: the pandemics within the pandemic. Clin Microbiol Infect, doi:10.1016/j.cmi.2021.05.022 (2021).

11 Zhu, X. et al. Cryo-electron microscopy structures of the N501Y SARS-CoV-2 spike protein in complex with ACE2 and 2 potent neutralizing antibodies. PLoS Biol 19, e3001237, doi:10.1371/journal.pbio.3001237 (2021).

12 Washington, N. L. et al. Emergence and rapid transmission of SARS-CoV-2 B.1.1.7 in the United States. Cell 184, 2587–2594 e2587, doi:10.1016/j.cell.2021.03.052 (2021).

13 Baum, A. et al. Antibody cocktail to SARS-CoV-2 spike protein prevents rapid mutational escape seen with individual antibodies. Science 369, 1014–1018, doi:10.1126/science.abd0831 (2020).

14 Weisblum, Y. et al. Escape from neutralizing antibodies by SARS-CoV-2 spike protein variants. Elife 9, doi:10.7554/eLife.61312 (2020).

15 Wang, P. et al. Antibody resistance of SARS-CoV-2 variants B.1.351 and B.1.1.7. Nature 593, 130–135, doi:10.1038/s41586-021-03398-2 (2021).

16 Kuzmina, A. et al. SARS-CoV-2 spike variants exhibit differential infectivity and neutralization resistance to convalescent or post-vaccination sera. Cell Host Microbe 29, 522–528 e522, doi:10.1016/j.chom.2021.03.008 (2021).

17 McCarthy, K. R. et al. Recurrent deletions in the SARS-CoV-2 spike glycoprotein drive antibody escape. Science 371, 1139–1142, doi:10.1126/science.abf6950 (2021).

18 McCray, P. B., Jr. et al. Lethal infection of K18-hACE2 mice infected with severe acute respiratory syndrome coronavirus. J Virol 81, 813–821, doi:10.1128/JVI.02012-06 (2007).

19 Winkler, E. S. et al. SARS-CoV-2 infection of human ACE2-transgenic mice causes severe lung inflammation and impaired function. Nat Immunol 21, 1327–1335, doi:10.1038/s41590-020-0778-2 (2020).

20 Moore, J. B. & June, C. H. Cytokine release syndrome in severe COVID-19. Science 368, 473–474, doi:10.1126/science.abb8925 (2020).

21 Herrmann, J., Mori, V., Bates, J. H. T. & Suki, B. Modeling lung perfusion abnormalities to explain early COVID-19 hypoxemia. Nature communications 11, 4883, doi:10.1038/s41467-020-18672-6 (2020).

22 Suh, Y. J. et al. Pulmonary Embolism and Deep Vein Thrombosis in COVID-19: A Systematic Review and Meta-Analysis. Radiology 298, E70–E80, doi:10.1148/radiol.2020203557 (2021).

23 Yao, Y. et al. D-dimer as a biomarker for disease severity and mortality in COVID-19 patients: a case control study. J Intensive Care 8, 49, doi:10.1186/s40560-020-00466-z (2020).

24 Kemp, S. A. et al. Recurrent emergence and transmission of a SARS-CoV-2 spike deletion H69/V70. bioRxiv, doi:https://doi.org/10.1101/2020.12.14.422555 (2021).

25 Peckham, H. et al. Male sex identified by global COVID-19 meta-analysis as a risk factor for death and ITU admission. Nature communications 11, 6317, doi:10.1038/s41467-020-19741-6 (2020).

26 Takahashi, T. et al. Sex differences in immune responses that underlie COVID-19 disease outcomes. Nature 588, 315–320, doi:10.1038/s41586-020-2700-3 (2020).

27 Pradhan, A. & Olsson, P. E. Sex differences in severity and mortality from COVID-19: are males more vulnerable? Biol Sex Differ 11, 53, doi:10.1186/s13293-020-00330-7 (2020).

28 Klein, S. L. et al. Sex, age, and hospitalization drive antibody responses in a COVID-19 convalescent plasma donor population. J Clin Invest 130, 6141–6150, doi:10.1172/JCI142004 (2020).

29 Synowiec, A., Szczepanski, A., Barreto-Duran, E., Lie, L. K. & Pyrc, K. Severe Acute Respiratory Syndrome Coronavirus 2 (SARS-CoV-2): a Systemic Infection. Clin Microbiol Rev 34, doi:10.1128/CMR.00133-20 (2021).

30 Li, D., Jin, M., Bao, P., Zhao, W. & Zhang, S. Clinical Characteristics and Results of Semen Tests Among Men With Coronavirus Disease 2019. JAMA Netw Open 3, e208292, doi:10.1001/jamanetworkopen.2020.8292 (2020).

31 Fenizia, C. et al. Analysis of SARS-CoV-2 vertical transmission during pregnancy. Nature communications 11, 5128, doi:10.1038/s41467-020-18933-4 (2020).

32 Funk, T. et al. Characteristics of SARS-CoV-2 variants of concern B.1.1.7, B.1.351 or P.1: data from seven EU/EEA countries, weeks 38/2020 to 10/2021. Euro Surveill 26, doi:10.2807/1560-7917.ES.2021.26.16.2100348 (2021).

33 Berlin, D. A., Gulick, R. M. & Martinez, F. J. Severe Covid-19. N Engl J Med 383, 2451–2460, doi:10.1056/NEJMcp2009575 (2020).

34 Blanco-Melo, D. et al. Imbalanced Host Response to SARS-CoV-2 Drives Development of COVID-19. Cell 181, 1036–1045 e1039, doi:10.1016/j.cell.2020.04.026 (2020).

35 Chen, X. et al. Detectable Serum Severe Acute Respiratory Syndrome Coronavirus 2 Viral Load (RNAemia) Is Closely Correlated With Drastically Elevated Interleukin 6 Level in Critically Ill Patients With Coronavirus Disease 2019. Clin Infect Dis 71, 1937–1942, doi:10.1093/cid/ciaa449 (2020).

36 Zhang, J. et al. Serum interleukin-6 is an indicator for severity in 901 patients with SARS-CoV-2 infection: a cohort study. J Transl Med 18, 406, doi:10.1186/s12967-020-02571-x (2020).

37 Zhou, Z. et al. Heightened Innate Immune Responses in the Respiratory Tract of COVID-19 Patients. Cell Host Microbe 27, 883–890 e882, doi:10.1016/j.chom.2020.04.017 (2020).

38 Donlan, A. N. et al. IL-13 is a driver of COVID-19 severity. medRxiv, doi:10.1101/2020.06.18.20134353 (2021).

39 Petrey, A. C. et al. Cytokine release syndrome in COVID-19: Innate immune, vascular, and platelet pathogenic factors differ in severity of disease and sex. J Leukoc Biol 109, 55–66, doi:10.1002/JLB.3COVA0820-410RRR (2021).

40 Hadjadj, J. et al. Impaired type I interferon activity and inflammatory responses in severe COVID-19 patients. Science 369, 718–724, doi:10.1126/science.abc6027 (2020).

41 Cummings, M. J. et al. Epidemiology, clinical course, and outcomes of critically ill adults with COVID-19 in New York City: a prospective cohort study. Lancet 395, 1763–1770, doi:10.1016/S0140-6736(20)31189-2 (2020).

42 Broggi, A. et al. Type III interferons disrupt the lung epithelial barrier upon viral recognition. Science 369, 706–712, doi:10.1126/science.abc3545 (2020).

43 Major, J. et al. Type I and III interferons disrupt lung epithelial repair during recovery from viral infection. Science 369, 712–717, doi:10.1126/science.abc2061 (2020).

44 Cameron, M. J. et al. Interferon-mediated immunopathological events are associated with atypical innate and adaptive immune responses in patients with severe acute respiratory syndrome. J Virol 81, 8692–8706, doi:10.1128/JVI.00527-07 (2007).

45 Channappanavar, R. et al. Dysregulated Type I Interferon and Inflammatory Monocyte-Macrophage Responses Cause Lethal Pneumonia in SARS-CoV-Infected Mice. Cell Host Microbe 19, 181–193, doi:10.1016/j.chom.2016.01.007 (2016).

46 Tobin, M. J., Laghi, F. & Jubran, A. Why COVID-19 Silent Hypoxemia Is Baffling to Physicians. Am J Respir Crit Care Med 202, 356–360, doi:10.1164/rccm.202006-2157CP (2020).

47 Xie, J. et al. Association Between Hypoxemia and Mortality in Patients With COVID-19. Mayo Clin Proc 95, 1138–1147, doi:10.1016/j.mayocp.2020.04.006 (2020).

48 Malas, M. B. et al. Thromboembolism risk of COVID-19 is high and associated with a higher risk of mortality: A systematic review and meta-analysis. EClinicalMedicine 29, 100639, doi:10.1016/j.eclinm.2020.100639 (2020).

49 Ackermann, M. et al. Pulmonary Vascular Endothelialitis, Thrombosis, and Angiogenesis in Covid-19. N Engl J Med 383, 120–128, doi:10.1056/NEJMoa2015432 (2020).

50 Zhang, L. et al. D-dimer levels on admission to predict in-hospital mortality in patients with Covid-19. J Thromb Haemost 18, 1324–1329, doi:10.1111/jth.14859 (2020).

51 Lee, M. H. et al. Microvascular Injury in the Brains of Patients with Covid-19. N Engl J Med 384, 481–483, doi:10.1056/NEJMc2033369 (2021).

52 Dan, J. M. et al. Immunological memory to SARS-CoV-2 assessed for up to 8 months after infection. Science 371, doi:10.1126/science.abf4063 (2021).

53 Turner, J. S. et al. SARS-CoV-2 infection induces long-lived bone marrow plasma cells in humans. Nature, doi:10.1038/s41586-021-03647-4 (2021).

54 Reynolds, C. J. et al. Prior SARS-CoV-2 infection rescues B and T cell responses to variants after first vaccine dose. Science, doi:10.1126/science.abh1282 (2021).

55 Kosikova, M. et al. Imprinting of Repeated Influenza A/H3 Exposures on Antibody Quantity and Antibody Quality: Implications for Seasonal Vaccine Strain Selection and Vaccine Performance. Clin Infect Dis 67, 1523–1532, doi:10.1093/cid/ciy327 (2018).

56 Wu, F. et al. A new coronavirus associated with human respiratory disease in China. Nature 579, 265–269, doi:10.1038/s41586-020-2008-3 (2020).

57 Xie, X. et al. An Infectious cDNA Clone of SARS-CoV-2. Cell Host Microbe 27, 841–848 e843, doi:10.1016/j.chom.2020.04.004 (2020).

